# RoCi - A Single Step Multi-Copy Integration System Based on Rolling-Circle Replication

**DOI:** 10.1101/2024.09.13.612835

**Authors:** Martzel Antsotegi-Uskola, Vasil D’Ambrosio, Zofia Dorota Jarczynska, Katherina García Vanegas, Martí Morera-Gómez, Xinhui Wang, Thomas Ostenfeld Larsen, Jean-Marie Mouillon, Uffe Hasbro Mortensen

## Abstract

Fungi are often used as cell factories for homologous and heterologous production of enzymes and metabolites. One strategy to obtain high yielding strains is to enhance the expression level of the gene(s) responsible for production of the product by inserting multiple copies of the gene-expression cassette. Typically, this is achieved by transforming non-homologous end-joining proficient strains with large amounts of a DNA vector, which randomly integrates in multiple copies at different loci, or more often, into a single locus with copies arranged as mixed orientation repeats. The majority of strains produced in this manner are unstable and substantial screening is necessary to identify strains with high and stable production. Moreover, the randomness of the insertion processes makes it difficult to determine how and where the copies are positioned in the genome. To this end, we envisioned that the instability of gene clusters made by the classical method is mostly due to the presence of a mixture of directly and inverted repeats. In such clusters, hairpins formed by inverted repeats may cause frequent recombinogenic lesions during replication to induce gene-expression cassette copy-loss by direct-repeat recombination. It is therefore possible that strains with gene-expression cassette clusters made solely by direct repeats would be more stable. Using *Aspergillus nidulans* as a model, we tested this idea and developed RoCi, a simple and efficient method to facilitate integration of multiple directly repeated gene-expression cassettes into a defined genomic locus through rolling-circle replication without pre-engineering requirements for strain preparation. In addition, we demonstrate that RoCi can be performed without *E. coli* based cloning, making it compatible with medium-high throughput experiments. Analyzing strains produced by RoCi, we have constructed strains bearing up to 68 *mRFP* GECs and we show that an *mRFP* multi-copy gene-array supports high and stable mRFP production for at least ∼150 generations on solid medium. In liquid culture we observed a minor average copy loss at 1 L scale. This loss could be eliminated by extending the gene-expression cassette with a crippled selection marker. To demonstrate the strength of the method, we used it to produce stable and high yielding cell factories for production of the specialized metabolite cordycepin on solid medium and of the enzyme β-glucuronidase in submerged culture. Finally, we show that RoCi can also be applied in the industrial workhorses *A. niger* and *A. oryzae* indicating that RoCi is generally applicable in fungi.

## Introduction

The saprophytic lifestyle of filamentous fungi requires the secretion of large repertoires of degradative enzymes and specialized metabolites (SMs) (1–3), and the superior secretory capacity of fungi in combination with their ability to propagate in low-cost culture media have made them a favorite choice for bioproduction of homologous and heterologous industrial enzymes and metabolites for the food and pharma industries (4–7). This choice is further substantiated by the fact that the fungal secretory pathway offers an efficient protein folding machinery coupled to folding quality control as well the potential to add post translational protein modifications (8,9); attributes which are lacking in bacterial production systems.

The popularity of cell factories based on filamentous fungi has sparked a substantial interest in developing efficient genetic tools for speedy strain construction. In this context, a big leap forward includes development of non-homologous end-joining (NHEJ) deficient strains and the implementation of CRISPR technologies, which together have set the stage for efficient gene targeting in filamentous fungi (10–15) using gene-targeting substrates based on PCR fragments and even oligonucleotides (16). Lately, it has been shown that even complex gene-targeting substrates can be assembled by *in vivo* by homologous recombination (HR) eliminating the need for *E. coli* cloning (17).

With cell factories in place, a popular and simple strategy to enhance production yields is to express multiple copies of the necessary production gene(s) (18,19). In baker’s yeast *Saccharomyces cerevisiae*, self-replicating 2µ based plasmids have been extensively used to express multiple copies of a <u>G</u>ene <u>E</u>xpression <u>C</u>assette (GEC) (Fang et al., 2011; Kim et al., 2014). For example, insulin is made in this manner (20–22). Inspired by this method, AMA1 based autonomous replicating plasmids have been developed and used in filamentous fungi (23,24), but they do not propagate in a highly stable manner and are to our knowledge not used in production.

More stable production strains can be achieved by integrating the GEC into a defined well-characterized genomic site (25,26). However, with this strategy only a single GEC is integrated into the genome, which limits GEC expression levels. To bypass this difficulty, multiple copies of a GEC cassette can be integrated into natural repetitive sequences in the genome. In *S. cerevisiae* sequences like rDNA, Ty and δ-sequences have been targeted for this purpose. Drawbacks of this strategy is the struggle to pinpoint the integration sites due to the hundreds of potential locations (27,28); and when mapped out, integrations tend to occur in only a handful of sites despite the abundance of possibilities (28–30). Moreover, the copy number of strains created in this manner tend to be unstable perhaps as the genomic repeats often exist in clusters (31–33). Multi-copy integrations in filamentous fungi have classically been achieved by transforming NHEJ proficient strains with large amounts of a circular DNA vector. In this setup, the DNA vector randomly integrates in multiple copies at different loci and/or into a single locus with copies arranged as direct- and inverted repeats (DRs and IRs). The drawback of this method is that they are difficult to characterize and that copy numbers typically decrease as the strain is propagated, requiring extensive screens to find stable strains (34–38).

Alternatively, stable multi-copy GEC based production can ideally be achieved by integrating numerous GECs individually into the genome at defined and well-characterized integration sites. This can be achieved by inserting a common landing platform for gene targeting at desirable loci in the genome, which can be subsequently targeted for gene amplification mediated by DNA DSBs introduced by e.g. I-SceI or CRISPR nucleases. This strategy has been successfully applied in baker’s yeast (39) and filamentous fungi (40). Although this strategy enables the construction of stable multi-copy strains, it requires significant work to establish the basic strain for gene amplification. Hence, it requires months of work to introduce the setup into a new species or into a new strain background and a simple- and versatile method for introducing multiple gene-copies into defined genomic sites to allow for high- and stable expression levels is desirable. Here, we present a straightforward and efficient method, RoCi, which allows multiple copies of a gene of interest to be integrated into a single defined locus creating a cluster of DRs based on the principle of rolling-circle replication. Moreover, we demonstrate that the resulting GEC gene-clusters allow for stable production suitable for at least 1 L scale fermentations and use the technology to dramatically enhance production of the SM cordycepin and a β-glucuronidase from *E. coli*.

## Materials & Methods

### Strains and media

All *Aspergillus* strains used and generated in this study are listed in Supplementary Table S2. All strains were cultivated in solid minimal medium (MM) (1 % glucose, 1x nitrate salt solution (41), 0.001% Thiamine, 1x trace metal solution (42) and 2 % agar). For bioreactor experiments, strains were cultivated in liquid MM with 2% glucose. All transformations were plated in solid transformation medium (TM), which shares the recipe with solid MM except for the carbon source. The glucose is replaced by 1 M sucrose. For β- glucuronidase experiments, strains were inoculated on YPD (10 g/L yeast extract and 20 g/L Peptone) 2 % glucose. All media were supplemented with 10 mM Uridine, 10 mM Uracil and/or 4 mM L-Arginine when necessary.

All plasmids were propagated using *Escherichia coli* DH5α solid (2% agar) and liquid LB medium with 100 mg/mL ampicillin.

### PCR and Plasmid Construction

PCR reactions were performed in 35 cycles using the proof-reading PfuX7 polymerase (43). All PCR reactions were made using the touch-down program starting at an annealing temperature of 68°C, lowering 0.5°C in each cycle. Standard reaction volumes were 50 µL containing 1x CXL Buffer (20 mM Tris/HCl pH 8.8, 10 mM KCl, 6 mM (NH_4_)_2_SO_4_, 2 mM MgSO4, 0.1 mg/mL BSA and 0.1% Triton X-100), 0.2 mM dNTPs, 0.4 μM primers (Integrated DNA Technologies (IDT), Belgium), 1 U PfuX7, 10-30 ng of gDNA or plasmid DNA. PCR reactions were purified using the Zymoclean^TM^ Gel DNA Recovery Kit (ZYMO RESEARCH). All primers used in this study are listed in Supplementary Table S3.

All generated plasmids were assembled by USER cloning (44). Cas9-CRISPR-tRNA vectors were constructed as previously described (16), using pFC330 as backbone (10). For plasmid pDIV088 assembly, pDIV068 was used as backbone (45), and plasmids pDIV1050 and pDIV1051 were assembled using pDIV131 as backbone (45). Finally, plasmid pDIV0941 was assembled using pU2002A as backbone (25). The GenElute Plasmid Miniprep Kit (Merck) was used for plasmid purification. All plasmids used and generated in this study are listed in Supplementary Table S1.

Linear repair templates for transformation were obtained by PCR amplification of the Gene Expression Cassette (GEC) from the plasmid, and the resulting fragment was gel purified using the Zymoclean^TM^ Gel DNA Recovery Kit (ZYMO RESEARCH) to avoid any carry over of template. The smaller size plasmids for transformation are generated by *in-vivo* assembly of two PCR fragments with overlapping regions as shown in Figure 4B & Supplementary Figure S9. Multiple PCR reactions of each fragment were pooled and purified by gel purification using the Zymoclean^TM^ Gel DNA Recovery Kit (ZYMO RESEARCH) and used for transformation.

### Fungal transformation, protoplastation and strain validation

Protoplastation was performed as shown by Nielsen et al., (46) and transformations were made following the method presented by Vanegas et al., (12). For each transformation 0.75 µg of CRISPR vector pDIV073 or pDIV1049 were used. Depending on the transformation, the repair template concentrations co-transformed with the CRISPR vector were: 0.75-1 µg for the linear repair template, 0.75 µg of the vector and 0.4 pmol of each fragment for the *in-vivo* assembly-based transformation. Transformation plates were incubated at 30°C, except for the *A. nidulans* transformation plates, which were incubated at 37°C. To select the strains with multiple *mRFP* copies, fluorescent colonies in solid MM plates were photographed using the setup described by Vanegas et al., (12) adjusting the exposure time to 0.5 s. The colonies that showed a higher fluorescence than the reference strain NID2699 (45) were selected for validation by Southern-blot. Fluorescence intensities were also assessed using the VILBER FUSION FX with the F-750 filter at super sensitivity.

### Genomic DNA extraction

Two main protocols for DNA extraction have been used in this study. For ddPCR analysis, gDNA was extracted from spores collected from colonies on solid plate or mycelium from bioreactor experiments using the Quick-DNA Fungal/Bacterial Miniprep Kit (ZYMO RESEARCH). For Southern-Blot, gDNA was extracted from mycelium. Spores from the selected colonies were cultivated in liquid MM + Arg + Ura + Uri overnight. The next day, mycelium was harvested using miracloth and samples were frozen dry overnight. Freeze dried samples were turned into powder using a steel bead (5 mm; Qiagen) and the TissueLyser LT (QIAGEN). Nucleic acid extraction was performed following the protocol described by Gautam, 2022, with some modifications.

### Southern-blot

The multi-copy strain validation was done by Southern blot. For DNA probe synthesis, the targeting site “B” of the DIVERSIFY gene landing platform was PCR amplified, up concentrated by isopropanol precipitation (48) and labeled using the ThermoScientific^TM^ Biotin DecaLabel DNA Labeling Kit. *A. nidulans* and *A. oryzae* samples were digested using the SalI (NEB, New England Biolabs) restriction endonuclease and the *A. niger* samples were treated with BamHI (NEB). A total of 4 µg of digested DNA were run from each sample in a 1% agarose (w/v) gel. DNA transfer from the agarose gel to the positively charged nylon membrane (Hybond-N+, Amershan Biosciences) was made by capillarity. For hybridization, a 1% (w/v) BSA, 1 mM EDTA, 0.5 M NaPO_4_ pH = 7.2 and 7% (w/v) SDS buffer was used (49) and 42°C as overnight hybridization temperature. For membrane treatment and development, the ThermoScientific^TM^ Biotin Chromogenic Detection Kit was used following manufacturer’s instructions.

### Copy Number Determination

The digital droplet PCR (ddPCR) for *mRFP* copy number quantification was performed following manufacturer’s instructions for the ddPCR Supermix for Probes (No dUTP) (BioRad, Cat#1863023). DNA digestion was performed by adding the restriction endonuclease MseI in the ddPCR mix and incubating the mix for 30 min at 37°C.

Then, droplets were generated using the BioRad Automated Droplet Generator and the PCR reaction was performed following specific thermal cycling conditions. Finally, the readout of the ddPCR was made using the BioRad QX200 Droplet Digital PCR System and the data analysis was done using the BioRad QuantaSoft analysis software, version 1.7.4.0917.

The probes were designed using IDT PrimeTime qPCR Probes online tool. The *mRFP* probe was labeled with 5′ 6-FAM/Zen/3′Iowa Black FQ. The *oliC* gene was used as reference gene for all organisms and the probes were labeled with 5′ HEX/Zen/3′IB FQ. All probes used in this study are listed in Supplementary Table S5.

### Inoculum Preparation and Cultivation conditions

Strains were cultivated in solid MM for spore harvesting, MM + Arg + Ura + Uri for *A. nidulans,* 4 days of incubation at 37°C, and MM + Ura + Uri for *A. niger* and *A. oryzae,* 6 days of incubation at 30°C. Spores were harvested with 5 mL of 0.02% tween 20 solution, filtered through miracloth and spun down at 10,000 g for 1 min and resuspended in 1 mL of 0.02% tween 20. Spore concentration was determined with a Thoma cell counting chamber.

Batch cultivations were performed in two biological replicates in 1.0-L Biostat Qplus bioreactors (Sartorius Stedim Biotech, Germany). The temperature was controlled at 37°C, and pH values ranging from 3 to 5 (when desired) were maintained by the automatic addition of 1 M NaOH/1 M H_2_SO_4_ and measured by pH sensors (Model EasyFerm Plus K8 160, Hamilton). The volumetric flow rate (aeration) values ranged between 0.1-1 vvm, and the stirring between 100-800 rpm according to the cultivation stage. The Bioreactor cultivation program was designed and performed by the fermentation software Lucullus®. The fermentation conditions are summarized in Supplementary Figure S1. The working volume in the vessel was 800 mL of 2% glucose MM with corresponding auxotrophies. The bioreactors were inoculated to a final concentration of 10^6^ spores/mL. Samples for dry weight, and copy number analysis were taken at 14, 18, 22, 26 and 38 hours of incubation.

For the β-glucuronidase activity experiment, 100 mL of 2% glucose YPD medium were inoculated to a final concentration of 10^6^ spores/mL in 500 mL shake flasks. Strains were cultivated for 24 h at 37°C, shaking at 150 rpm.

### Dry Weight Determination

0.45 μm PES filters (Frisenette) were dried in a microwave oven at 150 W for 20 min and weighed. The dried and weighed filters were used to filter 5 mL of culture in a vacuum filtration pump. The filters, now with the biomass, were dried in the microwave at 150 W for 20 min, put in a desiccator box for a day and then weighed.

### Stability assessment

For the solid media experiment, strain SDIV0526 with twenty-three *mRFP* GECs was inoculated on a big plate in triplicate and left in the incubator for a whole week until reaching a diameter of 10 cm. For the liquid media experiment, the selected strains (SDIV0526 & SDIV0527) were cultivated bioreactors in duplicates. Four samples were taken from the beginning to the end of the exponential phase and another one later in the stationary phase. DNA was extracted from each sample using the Quick-DNA Fungal/Bacterial Miniprep Kit (ZYMO RESEARCH) and the copy numbers of *mRFP* were analyzed by ddPCR, making changes in copy-number quantifiable and easy to follow.

### Generation number calculation of colonies

The number of nuclear divisions at the perimeter of a colony was estimated by the method suggested by Schoustra et al., (50). Envisioning a hypothetical hypha growing as a straight line, the authors used average numbers of 20 dividing nuclei in the apical cell and 13 µm spacing each nucleus as the basis to provide a conservative number of the nuclear divisions necessary to go from the center to the perimeter of the colony. In the mRFP production experiments with a colony radius of 3.95 cm, we estimate that nuclei at the perimeter are the result of ∼152 divisions. For the cordycepin pedigree colonies with a 1.5 cm colony radius, ∼57 divisions in each colony generation.

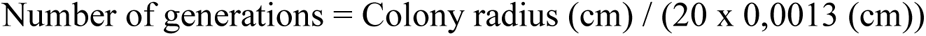

### Chemical analysis of strains

Strains were inoculated in solid MM and incubated at 37°C for 96 h. Metabolite extraction was performed as described by Smedsgaard (51). The analysis was performed using HPLC–DAD–HRMS on an Ultimate 3000 UHPLC system (Dionex, Sunnyvale, CA, USA) including a DAD detector, using the Waters ACQUITY UPLC HSS T3 (150 × 2.1 mm, 1.8 µm) at 40°C and a flow rate of 0.35 mL min−1. A linear water–MeCN gradient, containing 20 mM formic acid, was applied from 0% to 4% MeCN over 15 min, followed by an elution at 100% MeCN for 2 minutes, before returning to initial conditions. The samples were maintained at 4°C in the autosampler, and 1 µL was injected for each analysis. Mass spectrometric data was acquired on a MaXis HD QTOF-MS equipped with an electrospray ionization source. The mass spectrometer was operated in positive mode with a capillary voltage of 3500 V, and data recorded in a scan range from m/z 50 to 1400 at a rate of 10 scans/s. The drying gas flow rate was set to 11.0 L min−1, the temperature to 220°C and the nebulizer pressure was 2.0 bar. The collision cell settings included a transfer time 75 µs, collision cell RF 800 Vpp, and prepulse storage 8 µs. The mass spectrometer was calibrated using sodium formate. Data were evaluated with Bruker Compass DataAnalysis Version 5.3 (Build 556.396.6383).

### Quantitative Analysis of Cordycepin

A calibration curve for cordycepin was generated using a series of cordycepin standards (Sigma-Aldrich, United States) at concentrations of 0.10, 0.20, 0.39, 0.78, 1.56, 12.5 and 25 µg/mL. The yield of cordycepin in the samples was quantified by calculating the peak area from the extracted-ion chromatogram (EIC) corresponding to the [M + H]+ ion and comparing it to the calibration curve (Supplementary Figure S8B).

### β-glucuronidase activity assay

Fluorometric GUS Assay. For protein extraction, 100 mL of culture broth were filtered through Miracloth (Merck Millipore) and rinsed with miliQ water. The Miracloth was gently squeezed to remove excess liquid. Then, two small lab spoons (5 mm) of biomass were transferred to a 2 mL screw cap tube with a steel bead inside (5 mm; Qiagen) and frozen using liquid nitrogen. The samples were homogenized in TissueLyser LT (Qiagen) for 1 min at 45 Hz. 1 mL of GUS extraction buffer (52) was added, followed by homogenization for 2 min at 45 Hz. Samples were centrifuged for 3 min at 10 000g at 4 °C, and the liquid phase was collected by decanting into a new Eppendorf tube. The supernatant was centrifuged once more for 15 min at 10 000g at 4 °C. The supernatant was aliquoted and stored in −80°C. The total protein concentration was determined by standard Bradford assay (53).

The β-glucuronidase activity was determined according to the standard method (52) with slight modifications. 100 μL of substrate solution (4-MUG; Merck) was mixed with 95 μL of GUS extraction buffer and preheated for 2 min at 37 °C. Next, 5 μL of protein extract was added and such reaction assays were incubated for 5 min at 37 °C. Afterward, 5 μL of samples from reaction assays were withdrew, moved to the vials with 2 mL of stop solution, mixed and transferred into black 96-well microplate with a flat, transparent bottom (μClear; Greiner Bio-One). Fluorescence intensity was then measured on CLARIOstar^Plus^ microplate reader (BMG LABTECH) with excitation at 365 nm and emission at 455 nm. The plate with the samples was kept in the dark for 2−3 min prior to analysis to eliminate background luminescence Quantitation was performed using a six-level external calibration curve for 4-MU (Merck) with concentrations from 10 to 200 nM. The β-glucuronidase activity was normalized to total protein concentration. Statistical data analysis was performed using a two-tailed t-test with equal variances assumption and confidence interval of 99%.

## Results

### RoCi - A rolling-circle replication based scheme for incorporation of multiple directly repeated gene-copies into the genome

With the aim of developing a simple tool for construction of stable multi-copy GEC strains, we speculated that the production instability observed in strains made by transforming circular vector DNA into NHEJ proficient strains is mostly due to the formation of mixed repeat-type multi-gene clusters. Although copy loss in such cluster is mediated by direct repeat recombination, we envisioned that the presence of IRs is the main cause of recombination as they can form hairpins that are difficult to replicate. As a result, stalled- or collapsed replication forks can form (54–56), and these lesions are recombinogenic since repair of compromised replication forks may involve HR. Hence, in our working model, frequently formed lesions at IRs in mixed repeat clusters induce undesired DR recombination to cause GEC copy loss. Based on this rationale, we hypothesized that multi-gene clusters composed by direct repeats will be less prone to copy loss and perhaps allow for stable production in time frames fitting most laboratory experiments and possibly even in bigger production scales.

The task of building strains containing a cluster of directly repeated GECs in a defined genomic locus is not trivial, and no dedicated method for this task exists. To test the hypothesis, we therefore first conceived a scheme that potentially could be used for this purpose. As a starting point we exploited that specific DNA Double Strand Breaks (DSBs) produced by CRISPR- or meganucleases can be efficiently used to stimulate the integration of a linear <u>G</u>ene-<u>T</u>argeting <u>S</u>ubstrates (l-GTS) into a specific locus in a marker-free manner. Moreover, by performing the targeting experiment in NHEJ deficient strains, random polymerization of the l-GTSs by end-joining prior to integration is avoided, and only a single gene copy will therefore be introduced into the genomic DNA DSB, see Fig 1A (left). A GTS contained in a DNA circle (c-GTS) may also be applied for this purpose; and in this case, integration of a single gene-copy depends on gene-conversion or a double-crossover event, Fig 1A (right). These scenarios follow the classical synthesis dependent strand annealing (SDSA) and double strand break repair (DSBR) models that are usually used to explain DNA DSB repair involving sister chromatids or homologous chromosomes as repair templates (REFs-some reviews). Both SDSA and DSBR require that the 5’-ends of the DNA DSB are resected to produce large 3’ ssDNA overhangs; and, that one of these overhangs invades the repair template, see Figure 1B (left). At this stage, the 3’-end is extended by DNA polymerase to generate sequences that are complementary to the sequences of the other end of the DNA DSB setting the stage for second-end capture and insertion of the GTS into the site of the DNA DSB either by gene conversion or via a double crossover, Figure 1B (left). However, with c-GTSs, we envision a variant of this scenario where end-invasion results in rolling-circle replication prior to second-end capture. If so, gene-targeting using a c-GTS may deliver multiple copies, which per definition will be arranged as direct repeats, Figure 1B (right). Since this multi-copy integration system is based on <u>Ro</u>lling-<u>Ci</u>rcle replication, we will call it RoCi for convenience.

**Figure 1.**
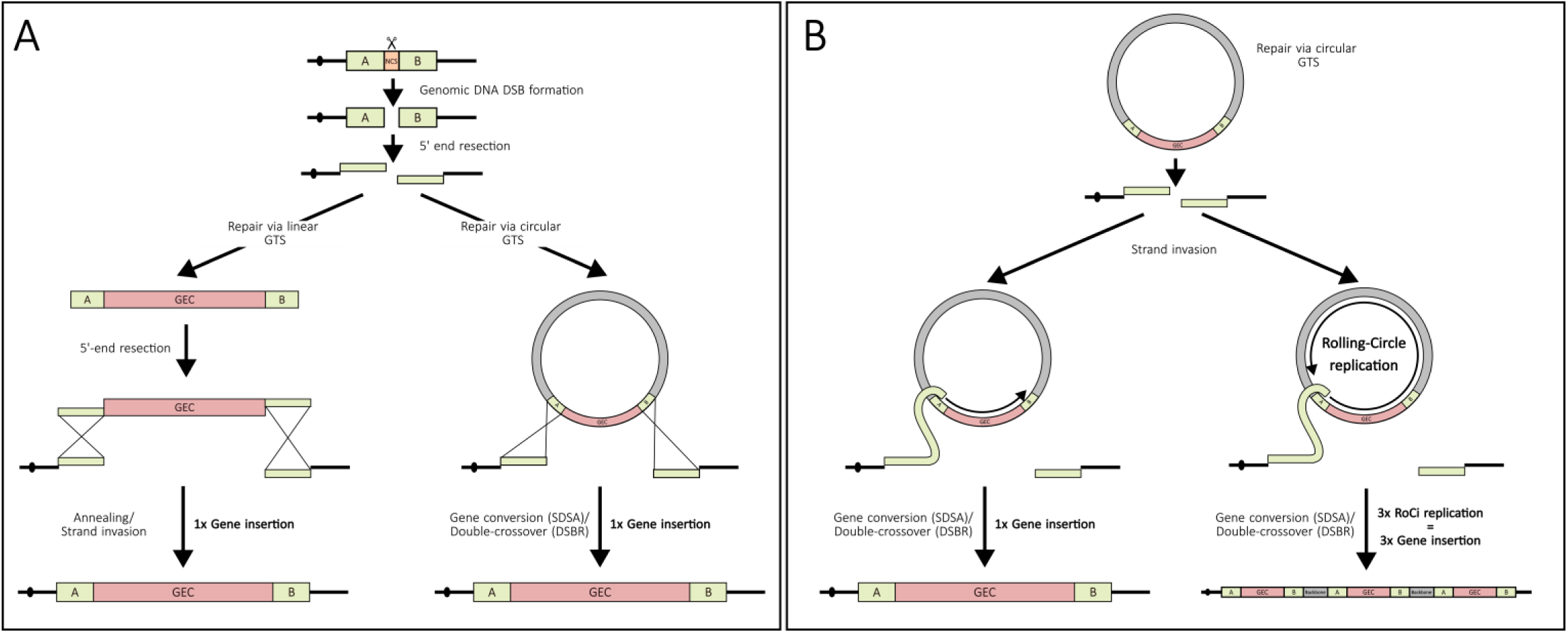
Clusters of directly repeated genes may be inserted into a genomic site by rolling-circle replication based gene targeting. Schemes showing the repair of a specific DNA DSB by using a l-GTS or a c-GTS as repair template (for details see main text). In both cases, the GTS is composed as a gene-expression cassette (GEI) flanked by the targeting sequences “A” and “B”. Gray box in the c-GTS is an *E. coli* vector backbone that may or may not be included in the circle. (**A**) Overview of key steps required to integrate a l-GTS (left) or a c-GTS (right) into a specific genomic DNA DSB induced by e.g. a CRISPR nuclease or a meganuclease. In the case of a l-GTS, the simplest integration scheme is initiated by 5’-resection of the ends of the DNA DSB and of the l-GTS. As a result, complementary ssDNA tails will be exposed at the ends of the DNA DSB and the l-GTS; and the matching ends can be joined by annealing to insert the l-GTS into the DNA DSB. In the case of the c-GTS, resection of the 5-ends of the DNA DSB sets the stage for integrating the cGTS into the genome as the result of SDSA or DSBR. (**B)** Integration of a cGTS is initiated when one of the two 5’-resected ends of the DNA DSB invades the circular GTS. Using the circle as a template for DNA replication, the invading strand is extended by DNA polymerase (three rounds of rolling circle replication is shown as an example). When DNA polymerase has replicated at least part of the region, which is complementary to the other end of the DNA DSB, second-end capture is possible and will proceed either via SDSA or DSBR. Importantly, the extension of the invading strand may be short to produce a single copy of the GTS (left); or long, if extension enters rolling-circle replication mode, to produce several copies of the GTS (right). Moreover, in the SDSA repair track, the extended and liberated 3’-end may re-invade the c-GTS for additional rounds of replication before completing repair (not shown).

### RoCi efficiently produces gene clusters of directly-repeated gene-expression cassettes in *Aspergillus nidulans*

To test whether RoCi is truly able to deliver a cluster of directly-repeated genes in a gene-insertion experiment, we took advantage of a gene-expression platform, DIVERSIFY, which we have previously developed (45). DIVERSIFY allows a gene-expression cassette (GEC) to be integrated into a <u>Co</u>mmon <u>S</u>ynthetic Gene Integration site, COSI site, which we have engineered into several different *Aspergilli* genomes at specific locations. Integration into a COSI site is stimulated by a Cas9/sgRNA complex that specifically cleaves in the COSI site; and proper targeting events can easily be detected in simple color based screens; e.g. our NHEJ deficient *A. nidulans* strain NID2696 (see Supplementary Table S2) contains the COSI site *COSI-1^uidA^*. Successful integration of a GEC into *COSI-1^uidA^* eliminates the *uidA* color-marker in the COSI sequence, and this event can therefore be identified in a simple blue-white screen (45). Using this setup, we co-transformed NID2696 with plasmid pDIV073, which encodes a CRISPR nuclease targeting *COSI-1^uidA^*, and either an l-GTS or an *E. coli* vector based c-GTS containing an *mRFP* (*PgpdA-mRFP-TtrpC*) GEC between the COSI targeting sequences “A” and “B”, see Figure 2A & B and see M & M. Transformants obtained with the *mRFP* l-GTS are expected to contain a single *mRFP* GEC (45); and in agreement with this view, essentially all transformants produced similar levels of red fluorescence (Figure 2A). In contrast, the red florescence produced by transformants obtained with the *mRFP* c-GTS varied in intensity. The lowest levels of fluorescence were similar to the levels observed with the transformants generated by *mRFP* l-GTS, and the highest levels were much brighter indicating that they contained multiple *mRFP* GECs Figure 2B.

**Figure 2.**
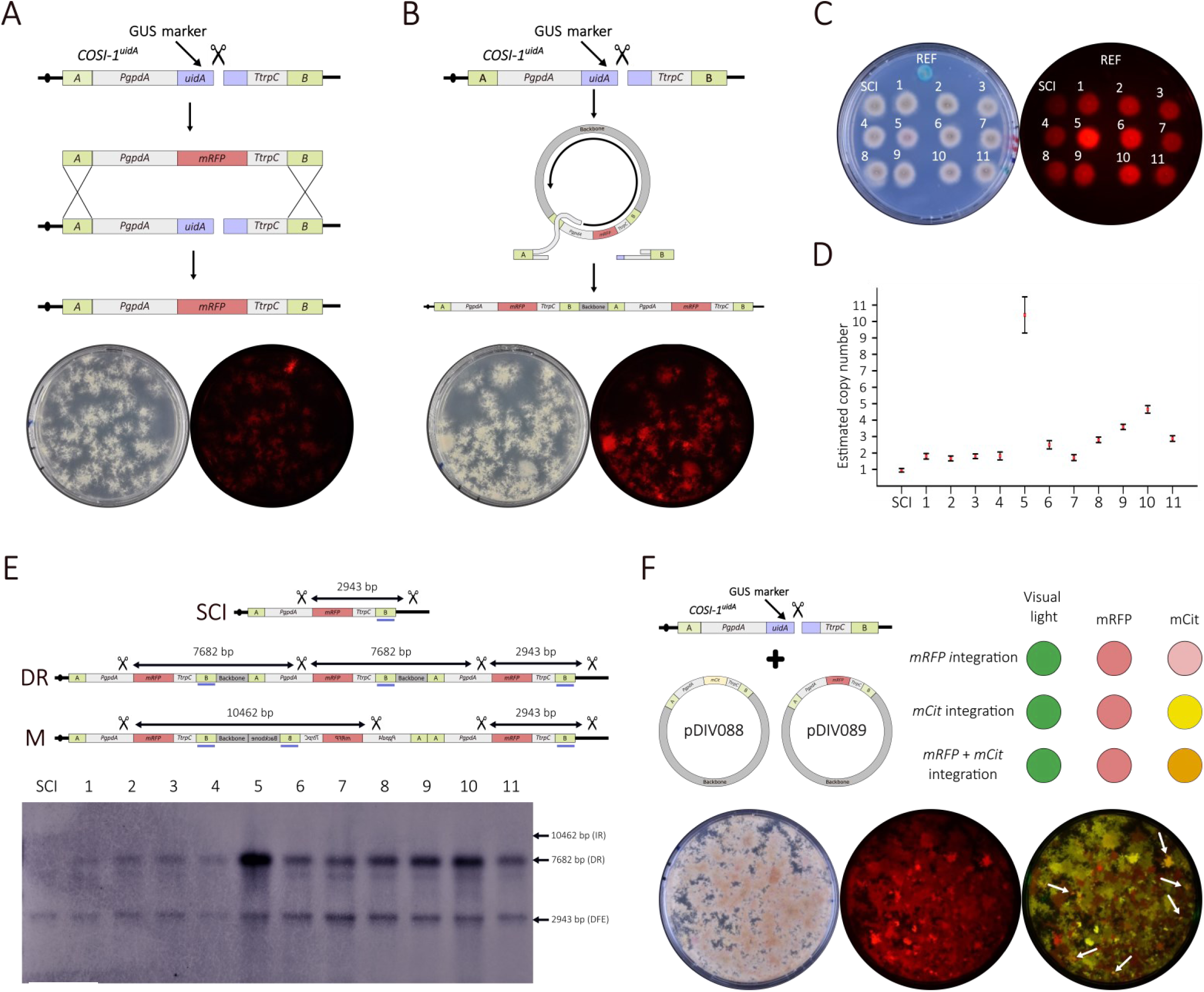
Rolling-circle replication based multi-copy integration in *A. nidulans*. (**A**) Scheme illustrating integration of a single *mRFP* GEC into a *COSI-1^uidA^* site (top). A specific CRISPR nuclease produces a DNA DSB in the *uidA* color marker indicated by a scissor. The break is repaired by HR using an l-GTS containing the *mRFP* GEC as the repair template. Images of transformants obtained by this scheme in visible-(left) and fluorescent light (right). (**B**) Scheme illustrating multi-copy integration of an *mRFP GEC* into a *COSI-1^uidA^* site by RoCi (top). A specific CRISPR nuclease produces a DNA DSB in *uidA*. The break is repaired by HR using a c-GTS containing the *mRFP GEC* as the repair template and via a process involving a rolling-circle replication step. Images of transformants obtained by this scheme in visible-(left) and fluorescent light (right). (**C**) Blue-white screening of RoCi transformants (stabs labelled 1-11) on solid MM-X-Gluc medium. Stabs of two control strains NID2696 (*COSI-1^uidA^*) and NID2699 (*COSI-1^mRFP^*) containing a <u>S</u>ingle *mRFP GEC* <u>C</u>opy Integration are labelled REF and SCI, respectively. Images obtained by visible-(left) and fluorescence light (right). (**D**) Determination of *mRFP GEC* copy numbers by ddPCR. Results from SCI and RoCi transformants 1-11. (**E**) Detection of the *mRFP GEC*s in SCI and RoCi transformants 1-11 by Southern-blotting. Graphic representations representing three different theoretical outcomes of *mRFP* GEC integration into *COSI-1* are shown in the top: SCI; an example of a cluster of directly repeated *mRFP* GECs (DR); and an example of a cluster of *mRFP* GEC repeats with mixed orientations (M). SalI cleavage sites in *COSI-1* and in *mRFP GEC*s are indicated by scissors and the expected sizes of the resulting fragments are indicated. The binding position(s) of the detection probe is(are) shown by a blue bar. Southern blot of SalI digested genomic DNA obtained from strain NID2699 (SCI) and RoCi transformants 1-11 are analyzed as indicated (bottom). Positions of fragments resulting from inverted repeats (IR), direct repeats (DR), and from the GEC at the <u>D</u>ownstream <u>F</u>lanking <u>E</u>nd of the COSI site (DFE) are indicated with labeled arrows. (**F**) Co-transformation of two c-GTS with two different fluorescent protein GECs can result in the integration of either one of the fluorescent protein GECs or, in some cases, in the integration of both. The three types of events are labeled with three different types of arrows as indicated.

To address whether the bright transformants obtained by RoCi contained at least one *mRFP* GEC in the COSI site, we selected and purified eleven transformants, which we judged to be more fluorescent by eye (Fig. 2B. For a full plate image, see Supplementary Figure S2). In a blue/white test, all selected transformants remained white indicating that the *COSI-1^uidA^*site was correctly targeted as expected and different fluorescence intensities could be appreciated between the different colonies, see Figure 2C.

We next assessed whether high levels of fluorescence in the selected transformants were due to the presence of multiple *mRFP* GEC copies in the same strain by ddPCR. For this analysis, we used the strain NID2699 (45), which contains a single copy of the *mRFP GEC* in the *COSI-1* site, as a control. These results demonstrated that the eleven transformants contained *mRFP GEC* copy numbers ranging from two to ten (Figure 2D). Importantly, we note that the deduced copy numbers of the strains correlated with the fluorescence intensities displayed by the strains (For full plate image, see Supplementary Figure S2).

We used Southern blotting to investigate whether the multiple GEC copies were inserted by RoCi. Specifically, we investigated whether the multiple *mRFP* GEC copies in the eleven transformants were all inserted into the *COSI-1* site in a head-to-tail fashion as expected from an integration mechanism involving a rolling-circle replication derived intermediate, see Figure 1B (right) and 2E. A comparison of the Southern-blot patterns obtained with the eleven transformants to the pattern generated by the *COSI-1^mRFP^* strain NID2699 demonstrated that in all cases only the *COSI-1* site has been targeted and that no copies were inserted as an inverted or aberrant repeat. Finally, the analysis showed that the intensity of the common band representing the individual direct repeats reflects the expected copy numbers determined by ddPCR. Altogether our analyses strongly indicate that the RoCi scheme can be used to insert multiple gene copies as direct repeats into a defined locus.

We further addressed whether multi-copy GEC insertion can be mediated by more complex RoCi integration events involving more than one strand invasion reaction. For example, after extension the liberated end could re-invade a c–GTS for further extension (Supplementary Figure S3A); or alternatively, the two ends of the DNA DSB could both invade a c-GTS and extend by rolling-circle replication, and the extended ends could then fuse by annealing to complete integration (Supplementary Figure S3B). To explore whether complex RoCi integrations occur with a significant frequency, we co-transformed strain NID2696 protoplasts with the CRISPR plasmid pDIV073 and two different c-GTSs, which contained the same backbone, but differed as one contains an *mCitrine* GEC (pDIV088) and the other an *mRFP* GEC (pDIV089), see Figure 2F. Ideally, if complex RoCi mediated integration involving both c-GTSs takes place, transformants displaying an orange phenotype should appear when they were analyzed for yellow fluorescence due to overlapping fluorescence between mRFP and mCit spectra. With this in mind, we have pointed out candidates that have integrated both GTSs on the transformation plate, see Figure 2F. To validate that these decisions reflect reality, we stabbed one colony of each type on a new plate and examined carefully for fluorescence and for the presence of the relevant genes by PCR, see Supplementary Figure S4, and these analyses confirmed our original phenotype scoring on the transformation plate. Together, our analyses demonstrate that complex repair may happen during a RoCi experiment.

### DNA circles assembled by *in vivo* recombination are efficient substrates for RoCi mediated multi-copy gene insertion

We have previously demonstrated that DNA circles can be efficiently assembled in filamentous fungi by fusing PCR fragments *in vivo* by homologous recombination (17), hence, eliminating time-consuming *E. coli* cloning steps and the need of an undesirable *E. coli* vector backbone. We therefore set out to explore whether DNA circles assembled by *in vivo* recombination could be used as c-GTSs in a RoCi experiment by testing whether an *mRFP* GEC could be inserted into a COSI site in this manner, Figure 3. Accordingly, we produced two PCR fragments that were able to fuse into a c-GTS containing an *mRFP* GEC. Two of the PCR fragment ends were able to fuse via 678 bp overlapping *mRFP* sequences and the other two ends via 50 bp assembly tags that were added to the relevant ends of the PCR fragments via primer tails, see Figure 3A. In parallel and as a control, we produced the corresponding bipartite l-GTS using PCR fragments that were not equipped with assembly tags.

**Fig. 3.**
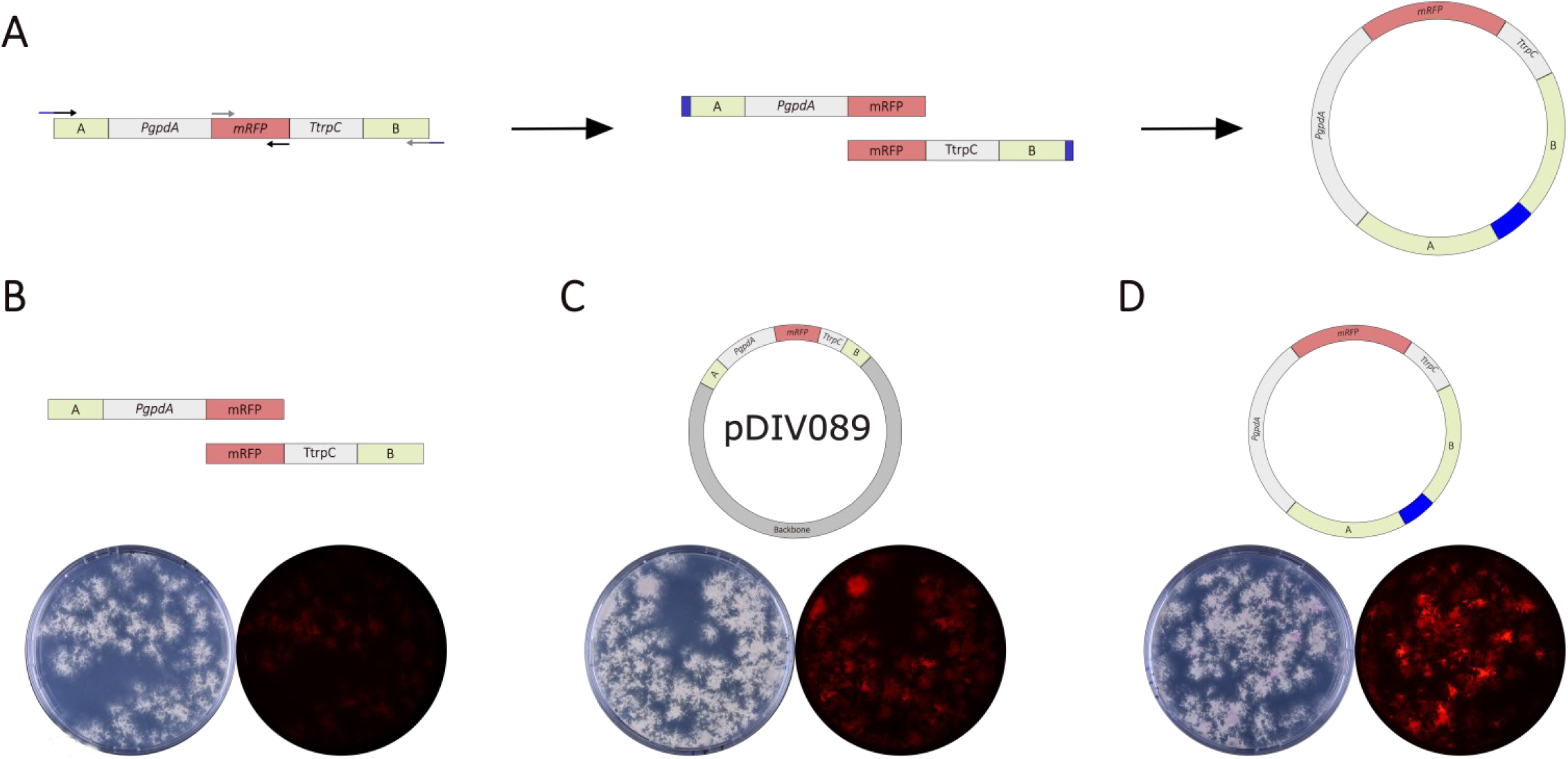
RoCi based multi-copy integration in *A. nidulans* using gene-targeting substrates made by *in vivo* assembly. (**A**) Generation of a c-GTS by *in vivo* assembly. First, two overlapping PCR fragments covering the entire GEC via the *mRFP* sequence are generated. The primers annealing to the “A” and “B” ends of the substrates are extended by assembly tags of identical sequences providing a second overlap to allow for c-GTS formation by *in vivo* assembly. (**B-D**) Transformation of the reference strain NID2696 with two PCR fragments that fuse into a bipartite l-GTS by *in vivo* assembly (**B**), a pre-formed c-GTS (**C**), and a c-GTS fused by *in vivo* assembly (**D**). All GTSs contain an *mRFP* GEC flanked by “A” and “B” targeting sequences matching the *COSI-1* site in *A. nidulans*. Schemes of the DNA parts are shown on the top of the panels. Transformation plates were imaged in visible light (left) and in fluorescent light (right).

Co-transformation of NID2696 protoplasts with the CRISPR plasmid pDIV073 and the bi-partite l-GTS, which cannot form a circle, are expected to produce transformants that contain a single copy of the GTS in the COSI site. In agreement with this view, the transformants emitted weak and almost uniform levels of fluorescence, see Figure 3B. We then examined the transformants obtained after co-transformation of *A. nidulans* protoplasts with pDIV073 and either pre-formed c-GTSs, containing an *E. coli* vector backbone DNA (Figure 3C), or the c-GTS made *in vivo* (Figure 3D). The number and ratio of non-fluorescent to fluorescent transformants in these two experiments were comparable strongly indicating that the bi-partite c-GTS PCR fragments were efficiently assembled into a functional c-GTS. Moreover, in both experiments, the individual transformants displayed different levels of fluorescence. These results are expected if the GTS is inserted into the COSI site via a process depending on a RoCi step where the GTS copy-number in the individual transformants should vary. By comparing the fluorescence levels of the transformants obtained with the c-GTS made by *in vivo* assembly to those obtained with the preassembled c-GTS, we find that the former appear stronger, see Figure 3D.

To examine this possibility in more detail, we did transformations in triplicate with a pre-formed c-GTS and with a c-GTS made *in vivo*. 3D fluorescence spectra of the transformation plates show that in the experiments made with a c-GTS made by *in vivo* assembly, there were more high peaks. All colonies from the transformation plates showing a significant peak in the fluorescence spectra were transferred to a new plate, one for each experiment (Supplementary Figure S5). The top five colonies with the highest peaks were selected from each plate and subjected to copy number analysis. The copy number distribution for the pre-formed c-GTS ranged from 2.63 to 17.5 copies and the median copy number was 4.3. For the c-GTS made by *in vivo* assembly the distribution ranged from 5.03 to 74 copies and the median copy number was 23, see Supplementary Table S6. Altogether, this set of experiments demonstrates that a c-GTS made by *in vivo* assembly acts as a highly efficient substrate for multi-GEC insertion via RoCi. Thus, all strains further along this study are generated using c-GTS made by *in vivo* assembly.

### Copy number and production stability of RoCi clusters in colonies and submerged culture

The fact that strains constructed by RoCi contain multiple GECs inserted as direct repeats raised the possibility that copies may be lost by direct-repeat recombination. To address this issue, we first investigated whether the rate of copy-loss influences overall GEC-copy numbers and GEC-production in a colony. Specifically, we selected the strain sDIV0526, which contains a considerable number of *mRFP* GECs in *COSI-1* (23 copies) for these analyses. On solid medium and in triplicate, we stabbed sDIV0526 in the center of a plate and allowed it to propagate to a diameter of ∼8 cm. At this stage, we examined the level of fluorescence at the perimeter of the colony by image analysis. Since, mitotic nuclear divisions in *A. nidulans* occur in the apical cells (57–59), this section of the colony contains the nuclei that are the products of the highest number of mitotic divisions in the colony; and thereby, most likely to suffer from copy loss events. Conveniently, fluorescence emitted by this section of the colony is easy to examine as the fluorescent signal is not compromised by spore formation. The image analysis showed that fluorescence at the perimeters of all three colonies appeared uniform and did not contain visible sectors/wedges with reduced fluorescence intensities (Figure 4A and for experiment in triplicate, see Supplementary Figure S6). Given the size of the colony, we roughly estimated that the nuclei at the perimeter of the colonies represent ∼150 mitotic divisions (see M & M) indicating that the cluster of *mRFP* GECs is quite stable.

**Figure 4.**
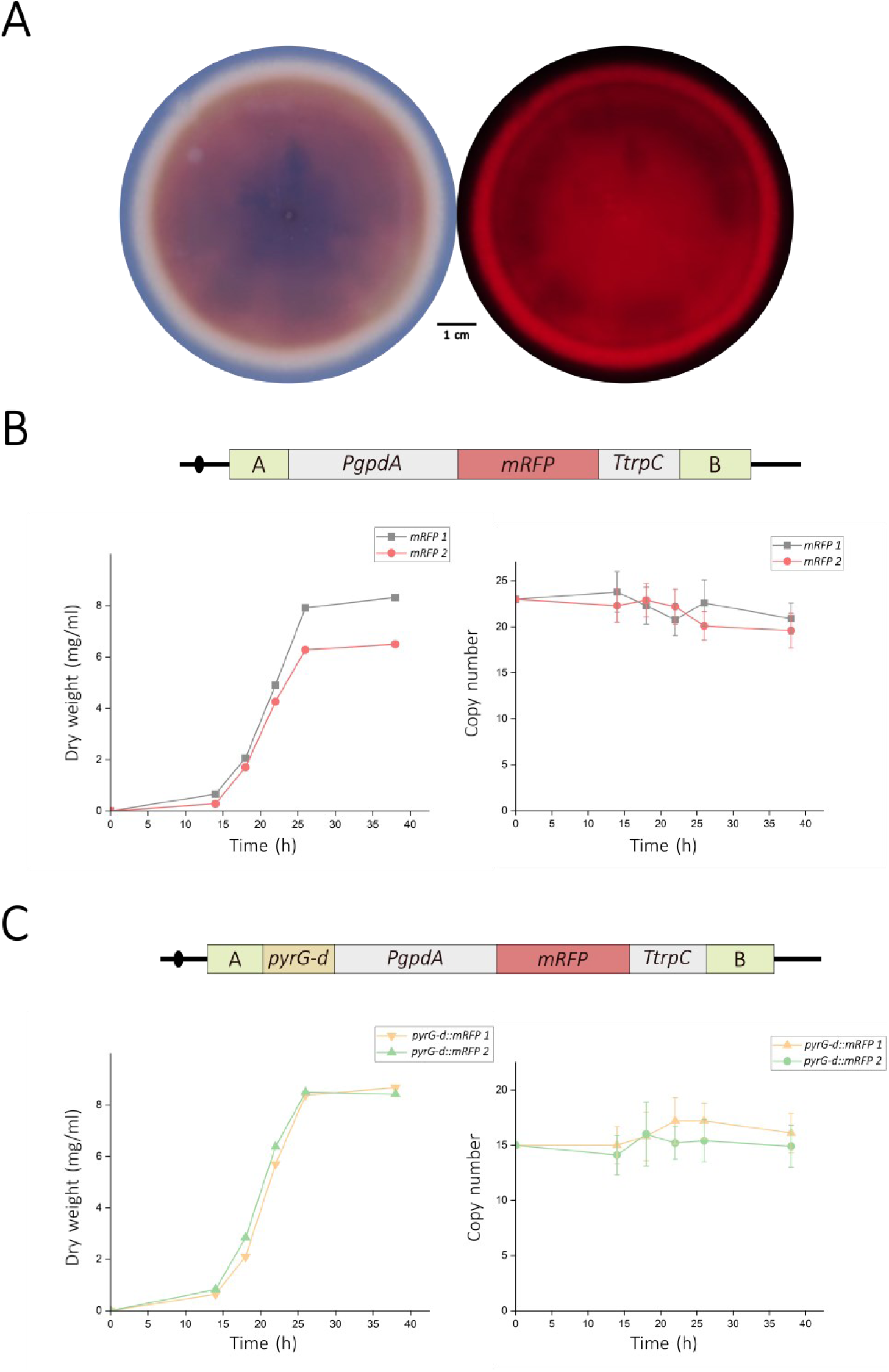
Directly repeated GEC cluster stability experiments in solid and submerged culture. (**A**) Colony formed by strain sDIV0526 containing 23 *mRFP* GECs. After 7 days of growth the colony was imaged in visible light (left) and in fluorescent light (right). (**B, C**) Biomass development and copy number development data of bioreactor cultivations in duplicate of strain sDIV0526 (**B**) and sDIV0527 (**C**) containing 23 *mRFP* GECs and 15 *pyrG-d::mRFP* GECs, respectively (C).

Fungal protein productions are mostly performed by using submerged cultures. To evaluate whether RoCi strains could be advantageously used for protein production, we therefore determined *mRFP* GEC copy-numbers by ddPCR throughout a growth experiment in 800 mL liquid MM 2% Glucose medium in a bioreactor in duplicate. As an inoculum we used spores harvested from five plates of strain sDIV0526. By performing ddPCR on DNA obtained from a fraction of these spores, we estimated that the spores contained an average copy number of 23 prior to the experiment. After inoculation, we followed growth for 40 hours where the cultures have reached stationary phase after 25 hours, see Figure 4B. This growth was accompanied by a modest reduction of 12% in *mRFP* GEC copy numbers in the two strains. Although this reduction may be acceptable in an experiment of this scale; it may raise a concern if the goal is to produce protein from larger volumes. To address this concern, we applied a classic strategy aiming at preserving copy numbers in clusters of repeated genes by including defective selectable marker genes in the repeats (60). We implemented this strategy by constructing RoCi strains where the *mRFP* GEC repeats were expanded with a *pyrG* gene controlled by a defective (truncated) promoter (*pyrG-d*) using a *pyrG*Δ strain (NID2696) as a starting point. Importantly, a *pyrG*Δ strain containing a single copy of the *pyrG-d*::*mRFP* GEC in *COSI-1* shows a significant growth delay, Supplementary Figure S7, showing that new strains suffering significant copy loss are unlikely to take over the population during growth. With this setup, we repeated the growth experiment in duplicate with strain containing *pyrG-d::mRFP* GEC in *COSI-1* (sDIV0527). The spore inoculum of this strain contained 15 GEC copies on average. Like in the previous experiment, growth was monitored for 40 hours, and the culture reached stationary phase after 25 hours. During the growth phase the growth rate of sDIV0527 was identical to the rate obtained by the *pyrGΔ* strain containing 23 copies of the *mRFP* GEC (sDIV0526) grown in Uridin and Uracil supplemented media, see Figure 4B & C. Indicating that the *pyrG-d::mRFP* GEC strain has sufficient copies to compensate for the poor *pyrG-d* promoter activity. Importantly, with the *pyrG-d::mRFP* GEC we did not observe any copy-loss during the entire experiment, see Figure 4C.

### RoCi can be advantageously used to enhance cordycepin production

Filamentous fungi are often used as cell factories for production of valuable metabolites, and we therefore decided to explore whether RoCi could be used to increase production of a specific SM. As a test case, we chose production of highly bioactive cordycepin (61–63), a nucleotide derived SM produced naturally by *A. nidulans* (64,65), but is most famously produced by *Cordyceps militaris* (66). We used *in vivo* assembly to produce a c-GTS containing the two relevant *A. nidulans* cordycepin production genes controlled by strong promoters and terminators (see M&M and Figure 5A) and RoCi to integrate the resulting cordycepin GEC into the *COSI-1^uidA^*site in multiple copies. In parallel, we also constructed a strain containing a single copy of the cordycepin GEC in this site. Colonies of the single-copy strain (sDIV0528) displayed a wild-type-like morphology on solid MM medium, Figure 5A. In contrast, the colonies obtained by RoCi displayed a reddish and crippled phenotype indicating toxic production levels of cordycepin, Figure 5A.

**Fig. 5.**
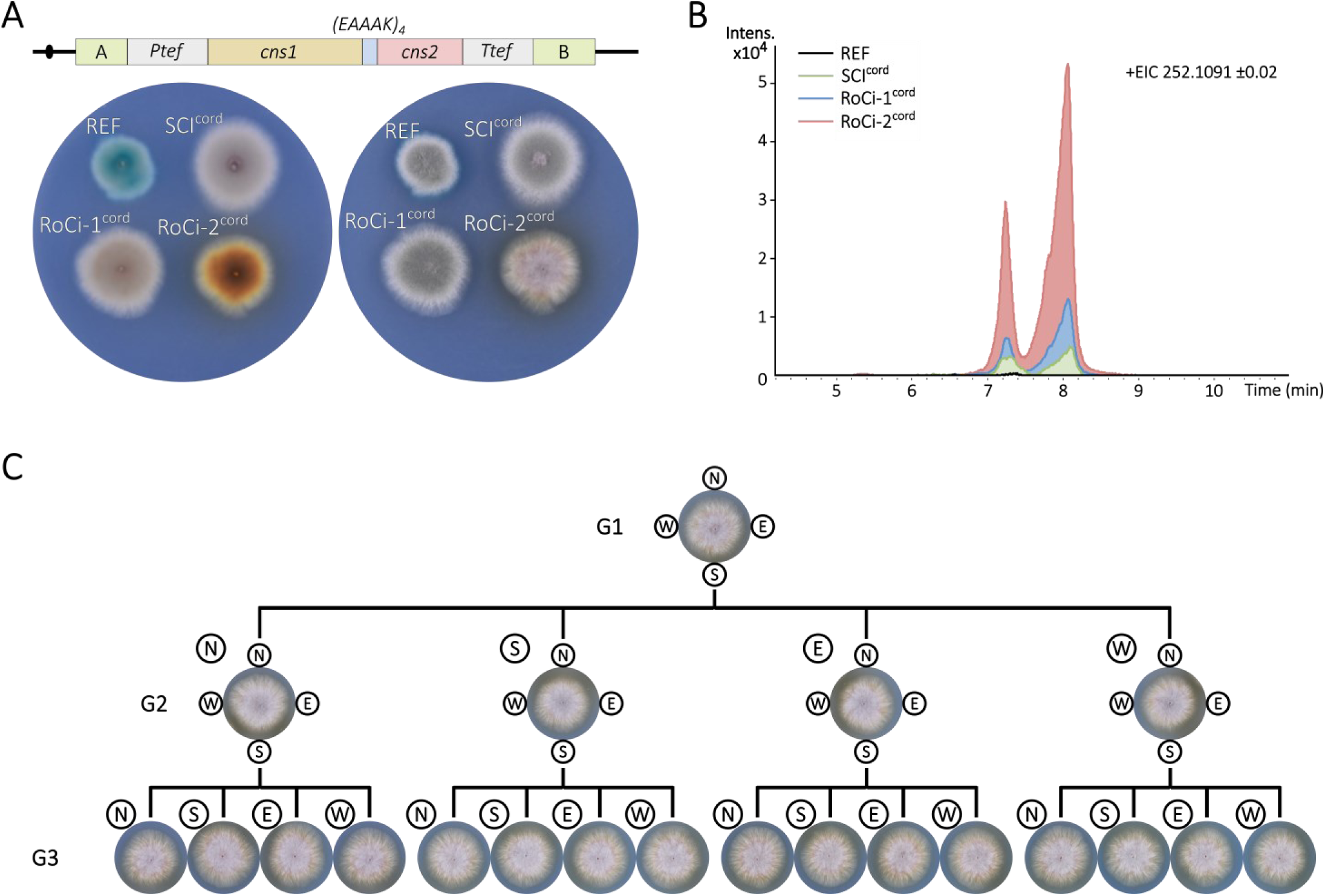
Cordycepin production GEC cluster is stable in colonies growing on solid. (**A**) Strains containing the *cns1(EAAAK)4cns2* GEC in different copy numbers were propagated on solid medium. Control strain NID2696 (REF); single cordycepin-GEC copy integration strain SCI^cord^ (sDIV0528); multi-copy strains Roci-1^cord^ & RoCi-2^cord^ (sDIV0529 and sDIV0530). (**B**) Extracted-ion chromatograms (EIC) of [M + H]^+^ of cordycepin in solid plug samples of the following strains: REF (black), single cordycepin-GEC (SCI^cord^, green), RoCi-1^cord^ (blue) and RoCi-2^cord^ (red). The extraction windows are for the exact m/z ±0.02, and the m/z value is the exact mass. (**C**) Colony generation pedigree of strain RoCi-2^cord^. Generation 1, 2, and 3 colonies were formed on solid MM medium as indicated and imaged after 96 h. At this time point, spores were collected at the perimeter at positions north, south, east and west (N, S, E and W circles), and stabbed on fresh solid MM medium to form the next generation colonies.

We next examined the colonies for intra- and extracellular cordycepin production by standard plug extraction, followed by HPLC/HRMS (M&M) to explore whether the crippled growth phenotypes correlated with cordycepin production, see Supplementary Figure S8. The reference strain produced barely detectable amounts of cordycepin. Importantly, the relative peak area of cordycepin [M+H]^+^ was considerably larger in extracts obtained from the strain with a single cordycepin GEC compared to the peak obtained with the reference strain, indicating a strong contribution from the cordycepin GEC. The single cordycepin-GEC strain (sDIV0528) produced a relative peak area corresponding to a concentration of 1.1 mg/mL. Next, we did the same analysis for two of the strains produced by RoCi, one with a weak (sDIV0529) and one with a strong phenotype (sDIV0530), see Figure 5A and Supplementary Figure S8A. Agreeing with the phenotypes, sDIV0529 and sDIV0530 produced 2.3 and 10 mg/mL of cordycepin corresponding to 2- and 9-fold higher production levels as compared the strain containing a single cordycepin GEC in *COSI-1* (sDIV0528), see Figure 5B.

The crippled growth phenotype observed for sDIV0530 strongly suggests that its cordycepin production levels are toxic. Hence, mycelia that have lost cordycepin GEC copies by direct repeat recombination should propagate faster and change morphology as the apparent fitness penalty of high cordycepin production is alleviated. To explore this possibility, we made a colony pedigree and examined whether the colony morphology would change after successive re-stabbing. Accordingly, we propagated sDIV0530 to a diameter of ∼ 3 cm. At this point, we isolated spores from four different positions in the perimeter of the colony; and used the material to make four-point re-stabs on a new plate containing solid MM medium, see Figure 5C. This procedure was repeated to create two generations of colonies. The sixteen colonies in the second generation all show a morphology like the founding colony; and none displayed sectors with a different morphology. Importantly, these observations strongly indicate that the cordycepin GEC cluster is stable during the experiment and that the ability to deliver desirable high production levels is maintained for many generations.

### A molecular copy-number counter can be used for simple identification of high-yielding cell-factory candidates after RoCi strain construction

Another major use of fungal cell factories is in enzyme production. Since GEC copy number is often key to high production yields, we investigated whether RoCi could be advantageously used for this purpose. As a model enzyme for heterologous production, we used β-glucuronidase encoded by *uidA* as production yields of this enzyme are easy to quantify. As a reference strain we used (sDIV0531) harboring the *COSI-1^mCit^* targeting platform instead of *COSI-1^uidA^*, see Figure 6A and Supplementary Table S2. sDIV0531 derived transformants with no GTS integration in the *COSI-1* site display yellow fluorescence, whereas correctly targeted transformants are identified by loss of yellow fluorescence due to the replacement of the *mCitrine* gene in the *COSI-1^mCit^* targeting site by the GTS. As a modification of the previous RoCi setup, we included a 2 kb *mRFP* GEC in the backbone of the c-GTS where it serves as molecular copy-number counter allowing for simple evaluation of the GEC copy-number after a RoCi experiment. This feature is useful in cases where the product itself cannot readily be detected by examining a simple phenotype displayed by the transformants. The *mRFP* GEC is not part of the targeting cassette of the c-GTS, therefore, transformants containing only one GEC copy can be scored as non-fluorescent. In contrast, transformants containing more than one copy emit red fluorescence as RoCi incorporates the entire c-GTS including *mRFP*, see Figure 6A. Lastly, the relative GEC copy number in the individual transformants can be quickly assessed by comparing their levels of red fluorescence as increasing c-GTS copy numbers in *COSI-1* are accompanied by increasing levels of red fluorescence.

**Fig. 6:**
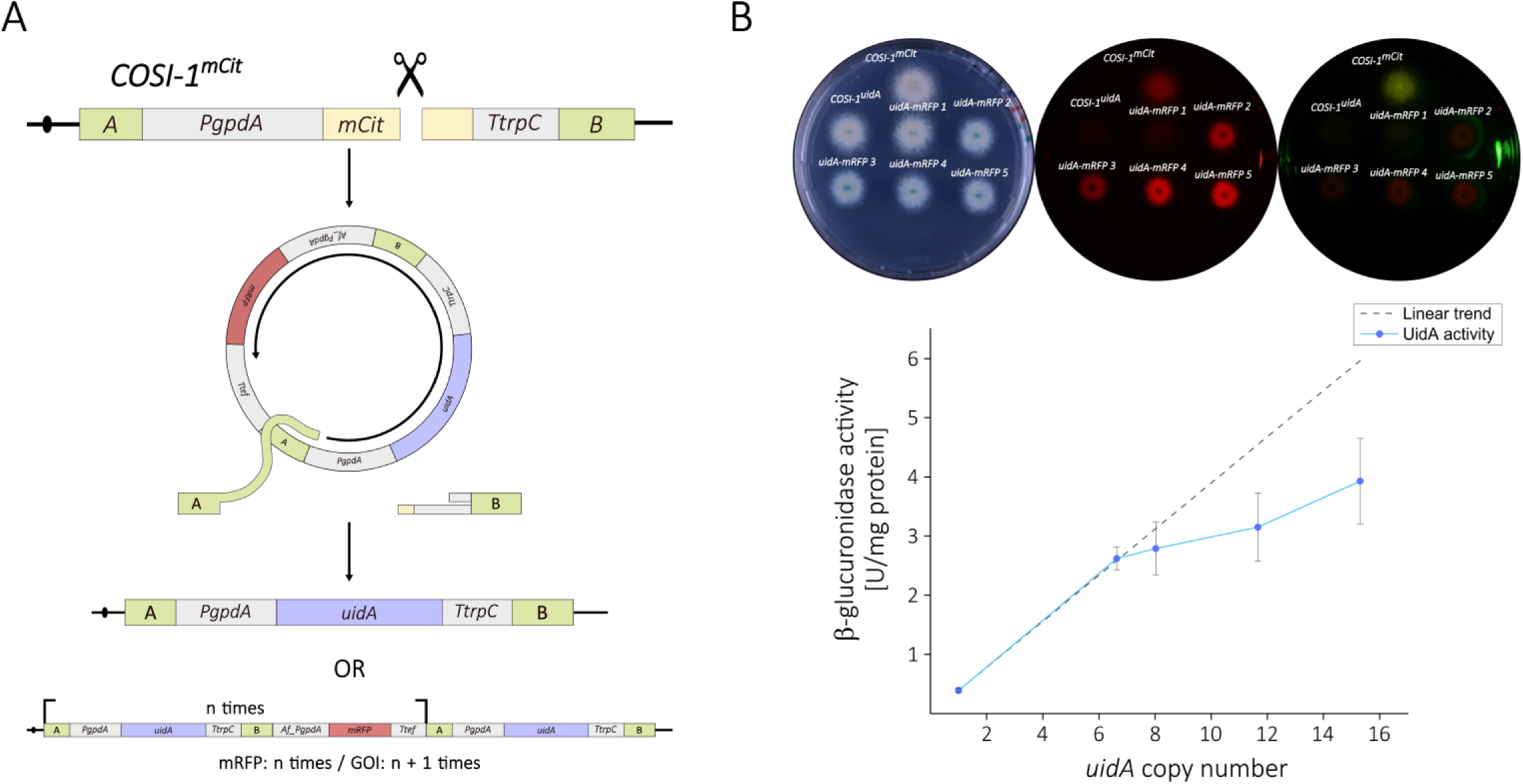
Multi-copy integration of *uidA* by RoCi. (**A**) Scheme illustrating multi-copy integration of the *in vivo* assembled c-GTS into a *COSI-1^mCit^* site by RoCi. A specific CRISPR nuclease produces a DNA DSB in *mCit*. The break is repaired by HR using a c-GTS containing the *uidA* GEC and the *mRFP* GEC as the repair template and via a process involving a rolling-circle replication step. This will result either in the integration of a single copy of the *uidA* GEC, as the repair will happen through the homology regions A or B, or in the integration of multiple copies of both the *uidA* GEC and the *mRFP* GEC; however, there will always be one more copy of the *uidA* GEC between the homology regions A and B. (**B**) Blue-white screening of strains sDIV0531 (*COSI-1^mCit^*) used for transformation, strain NID2696 (*COSI-1^uidA^*) as a single *uidA* copy control, and the RoCi transformants (*uidA-mRFP* 1-5) on solid TM-X-Gluc medium imaged in visible light (left), in red-(middle), and in yellow fluorescent light (right). (**C**) Graphic representation of ideal β-glucuronidase activity lineally increasing with the *uidA* GEC copy number relative to the level obtained with NID2696 (dashed line) and the real β-glucuronidase activity progression (blue line).

To test this setup, we co-transformed *A. nidulans* protoplasts of strain sDIV0531 with the CRISPR vector pDIV1049, which mediates insertion of the GTS into *COSI-1^mCit^*, and two PCR fragments that are able to fuse by *in vivo* HR to form a c-GTS containing *mRFP* and the *uidA* GEC (Supplementary Figure S9). To assess the use of the *mRFP* marker as a molecular copy-number counter that can be used to identify high-yielding cell-factories after a RoCi experiment, we determined *mRFP* GEC copy number and β-glucuronidase activity of selected transformants. Specifically, five colonies with no yellow fluorescence and with different red fluorescence intensities were selected from the transformation plate: a colony with no red fluorescence *uidA-mRFP* 1 (sDIV0532), two colonies with medium-low red fluorescence intensity *uidA-mRFP* 2 & *uidA-mRFP* 3 (sDIV0533 & sDIV0534), and the two colonies displaying the highest level of red fluorescence *uidA-mRFP* 4 & *uidA-mRFP* 5 (sDIV0535 & sDIV0536), see Figure 6B. As a control for the experiment, we added strain NID2696 since it harbors a single copy of *uidA* in *COSI-1* and no fluorescence markers.

As expected, the strain with no red fluorescence, *uidA-mRFP* 1, showed levels of β-glucuronidase activity comparable to those obtained with the control strain NID2696 that contains a single copy of *uidA* in the *COSI-1* site (*COSI-1^uidA^*). This result indicates that our new setup is able to identify a strain harboring a single *uidA* GEC copy. This conclusion was substantiated by ddPCR, which did not detect any *mRFP* copies (Table 1).

**Table 1.** *mRFP* copy number versus β-glucuronidase activity. (**) indicates significantly different production (p < 0.01) of GUS relative to the level obtained with NID2696.

| Strain | *mRFP* copy number | Estimated *uidA* GEC copy number | β-glucuronidase activity (U/mg protein) | Fold increase in β-glucuronidase activity |
| --- | --- | --- | --- | --- |
| NID2696 | 0 | 1 | $0,39 \pm 0,036$ | - |
| *uidA-mRFP* 1 | 0 | 1 | $0,32 \pm 0,046$ | 0,81 |
| *uidA-mRFP* 2 | $5,63 \pm 0,33$ | 6,63 | $2,62 \pm 0,195^{**}$ | 6,7 |
| *uidA-mRFP* 3 | $7,02 \pm 0,22$ | 8,02 | $2,79 \pm 0,443^{**}$ | 7,2 |
| *uidA-mRFP* 4 | $10,66 \pm 0,37$ | 11,66 | $3,15 \pm 0,576^{**}$ | 8,1 |
| *uidA-mRFP* 5 | $14,3 \pm 0,7$ | 15,3 | $3,93 \pm 0,721^{**}$ | 10,1 |

The remaining four strains, *uidA-mRFP* 2-5, all emitted red fluorescence, but with different intensities. Based on our visual analysis, the transformants were ordered according to increasing red fluorescence. This order correlated with increasing *mRFP* copy numbers as determined by ddPCR (Table 1). This analysis also showed that the *mRFP* copy numbers varied from six to fourteen indicating that they contain from seven to fifteen *uidA* GECs. Importantly, the increasing *uidA* GEC copy numbers were accompanied by increasing β-glucuronidase activity and they produced from seven- to ten-fold higher β-glucuronidase activity as compared to the single *uidA* GEC copy strain NID2696. We notice that the β-glucuronidase activity gain relative to the *uidA* GEC copy number is not entirely linear as it appears to level off at high *uidA* GEC copy numbers see Figure 6C & Table 1. Altogether, our experiments validate that this version of RoCi can be used to swiftly identify high GEC copy transformants. Moreover, it can serve as a useful tool to identify the best candidate cell factories for efficient heterologous enzyme production, in this case β-glucuronidase from *E. coli*.

### RoCi can be efficiently applied in other filamentous fungi

In a final set of experiments, we investigated whether RoCi could be used to insert multiple directly repeated GECs into specific genomic sites in other *Aspergilli*. To this end, we exploited previously constructed NHEJ deficient industrially relevant strains of *A. oryzae* (ORY7) and *A. niger* (NIG159) possessing the *COSI-1^uidA^* site (45). Both strains were used for RoCi experiments using the same CRISPR vector and the same *mRFP* GEC GTSs that we used above to target *A. nidulans*. Like with *A. nidulans*, co-transformation of *A. oryzae* and *A. niger* with the CRISPR vector and the l-GTS containing the *mRFP* GEC, produced transformants displaying low, but homogenous levels of fluorescence; in contrast, transformants obtained with the *mRFP* GEC in a pre-formed c-GTS produced fluorescence with variable intensity ranging from levels that were similar to those of the transformants obtained with the corresponding l-GTS to substantially higher levels (Figure 7A & B). For both species, we selected two transformants displaying high levels of fluorescence and demonstrated by blue-white screening, that the targeting event happened in the *COSI-1^uidA^* site (Figure 7C), and by ddPCR that they contained multiple *mRFP* GEC copies (Figure 7D). Moreover, Southern blot analyses demonstrated that the *mRFP* GEC copies in both transformants and for both species were organized as direct repeats in the *COSI-1* sites in agreement with an integration mechanism mediated by RoCi (Figure 7E). Hence, the RoCi principle also appears to apply to other filamentous fungi.

**Figure 7.**
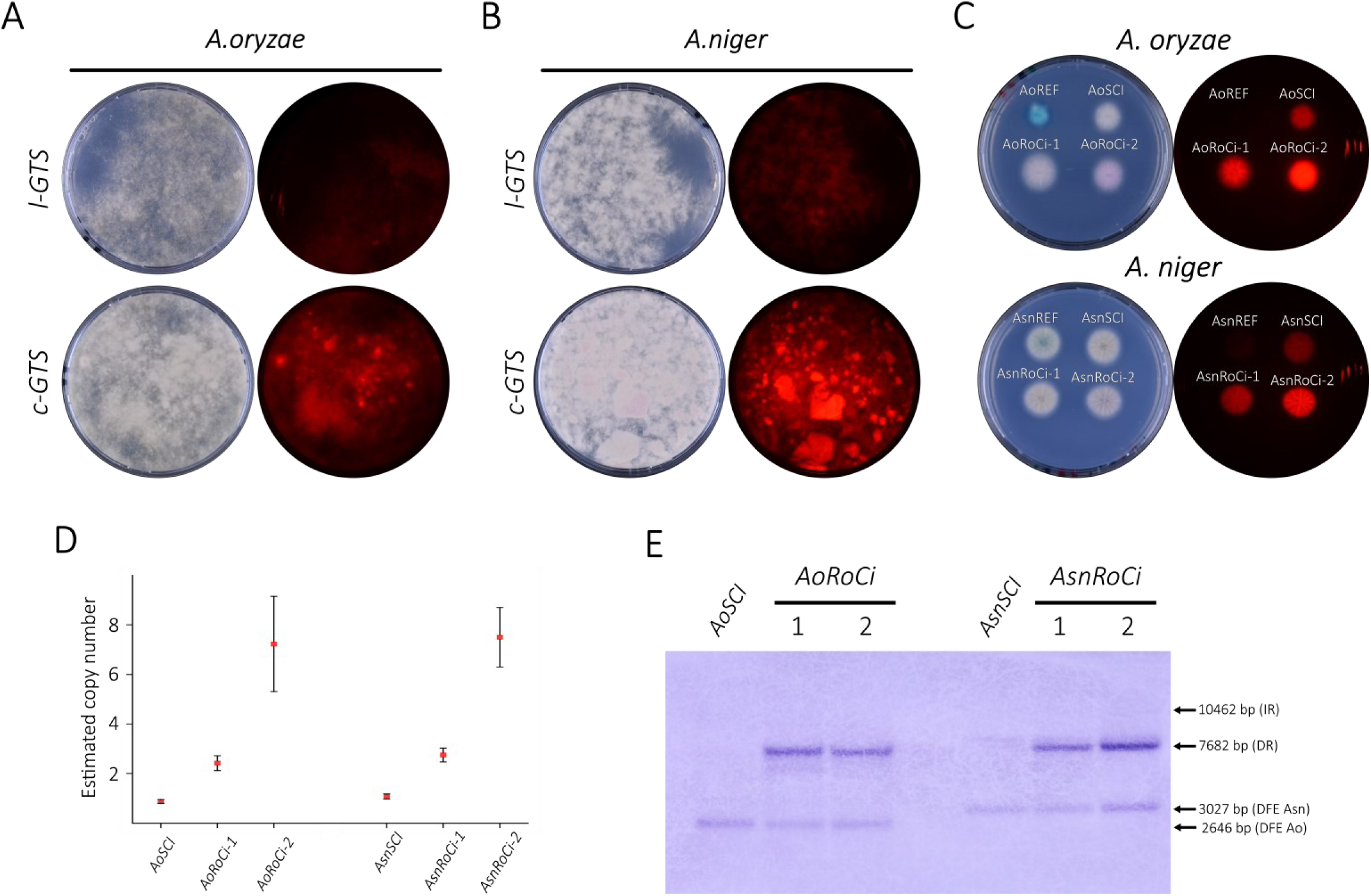
Rolling-circle replication based multi-copy integration in *A. oryzae and A. niger*. (**A, B**) *COSI-1^uidA^* sites in *A. oryzae* (**A**) *and A. niger* (**B**) were targeted with an l-GTS and a c-GTS containing an *mRFP GEC*. Transformations plates were imaged in visible light (left) and in fluorescent light (right). (**C**) Blue-white screening plates of RoCi transformants on solid MM-X-Gluc medium. Two control strains were stabbed, the reference strain used for transformation ORY7 (Ao*COSI-1^uidA^*), labelled as AoREF and a single-copy integration strain sDIV0537 obtained from the l-GTS transformation plate containing a single *mRFP GEC* copy integration, labelled as AoSCI (Asn*COSI-1^mRFP^*). Finally, two multi-copy strains, sDIV0538 and sDIV0539 (AoRoCi-1 and AoRoCi-2, respectively), obtained from the c-GTS transformation plates were stabbed. The corresponding master plate was made in the same way for for *A. niger* (Asn abbreviation). (**D**) Determination of *mRFP GEC* copy numbers by ddPCR. Results obtained with the *A. oryzae mRFP GEC* single copy strain AoSCI and two *A. oryzae* transformants obtained by RoCi; and from the *A. niger mRFP GEC* single copy strain AsnSCI and two *A. niger* transformants obtained by RoCi as indicated. (**E**) Southern blot of SalI digested genomic DNA obtained from the *A. oryzae* strain AoSCI and two *A. oryzae* transformants obtained by RoCi; and of BamHI digested genomic DNA from the *A. niger* strain AsnSCI and two *A. niger* transformants obtained by RoCi as indicated. Positions of fragments resulting from inverted repeats (IR), direct repeats (DR), and from the GEC at the <u>d</u>ownstream flanking <u>e</u>nd of the COSI site (DFE) are indicated with labeled arrows.

## Discussion

In this study, we show that repair of a genomic DNA DSB from a circle may involve a rolling-circle replication step. Hence, in addition to the expected one-copy insert, large clusters of directly repeated GTSs may be incorporated into the repaired site. Based on this observation, we have developed a novel method named RoCi that allows for simple construction of strains for overexpression of desirable genes. In this study we have used the DIVERSIFY strains bearing a COSI site to validate the method, but a synthetic insertion site is not a requirement to apply the method. This method can be used in any chosen insertion site without any pre-engineering of the strain as it only requires two targeting sites in the flanking regions of the GEC. Since all copies are integrated as direct repeats into a defined genomic locus, the genotype of the resulting strains can be quickly assessed by e.g. ddPCR. As an alternative, we have also devised a version of RoCi where the copy number can be assessed by including a visible marker in the circular vector-backbone containing the GTS. With the latter, strains with the highest copy-numbers can quickly be identified in an almost labor-free manner. To allow for *E. coli* cloning-free RoCi experiments based on c-GTS containing only sequences necessary for gene-expression, we further developed RoCi to include c-GTS that are assembled by *in vivo* recombination. Using RoCi based on c-GTS assembled by in vivo HR, we have constructed a strain with 68 *mRFP* GECs. In these experiments, large arrays of direct repeats can theoretically also be formed by inserting a long linear concatemer of GECs, formed by joining many individual parts in multiple successive annealing steps, which then inserts into the target site by HR. The result of this integration reaction will be the same as a RoCi reaction (Figure 1B, right). However, we strongly favor that the multi-gene integrations we observe occur by RoCi. Specifically, circle formation requires that the two DNA fragments are joined by a single 2^nd^-order fusion followed by a single 1^st^ order fusion as compared to the multiple 2^nd^-order fusions required to form a linear concatemer containing multiple GECs and where each fusion-step is expected to be slow due to the low concentration of each fragment in the nucleus. Lastly, we note that the highest copy numbers were obtained with the smallest c-GTS. Intuitively, one may envision that the number of replication cycles is higher with a small circle relative to with a large circle if the processivity of the DNA polymerase is the same. If so, this possibility may advantageously be considered in the planning of future RoCi experiments. At this point, we stress that the different c-GTSs used in this study are quite different and do not set the stage for fair comparisons to assess the impact of size on copy-number.

Importantly, the possibility of inserting many GECs into a specific integration site sets the stage for the construction of high-yielding cell factories. To this end, we tested whether RoCi could be used to make more efficient cell factories for SM and enzyme production and demonstrated that we could easily make strains producing 9-fold more cordycepin and 10-fold more UidA as compared to the yields obtained with strains containing only a single GEC copy.

The fact that RoCi produces clusters composed by direct repeats raises the questions whether unavoidable direct-repeat recombination would produce strains that are genetically unstable due to copy loss. If this process is frequent, the production advantages observed with the initial strains may be quickly lost, thereby limiting the use of RoCi strains dramatically. We have followed production in RoCi strains producing mRFP, and cordycepin and we find that production appears surprisingly stable. In particular, we were impressed by the fact that strains producing levels of cordycepin causing a growth defect phenotype, maintained this phenotype in three generations of colonies. In this context we note that the material used to initiate the final generation colonies had undergone approximately 100 nuclear divisions before it was stabbed onto the final plate. However, we stress that for products causing a larger fitness defect this may not be the case. More quantitatively, we noted that strains producing mRFP in liquid 800 mL cultures, lost 12 % of the GEC copies during 38 hours of cultivation starting from spores. Altogether, in our experiments copy loss does not appear to majorly impact production stability indicating that the GECs used in this study do not cause replication stress. However, this may not be true for other GECs, and it may be advisable to examine the sequences involved in RoCi for the presence of sequences that are known to cause replication stress like inverted-repeats, G-quadruplexes, and AT-rich regions (67).

If copy loss is a concern, we have shown that stability of RoCi strains can be enhanced by including a defective selectable marker in the c-GTS (60). In this case, mycelia that lost c-GTS copies will grow slower due to marker loss. For how long strains with this feature can propagate without marker-loss remains to be determined. In this context, we note that copy-loss is not the only result of HR in a cluster and that defective selectable markers have been used to select strains where the copy-numbers of gene-clusters have been further expanded (28,29).

Although RoCi strains may not be the choice for large scale industrial production, we envision that RoCi strains can be used in academic settings or in industrial pilot scale to deliver sufficient proof of principle to motivate time-consuming construction of more stable production strains. As RoCi strains can be made via c-GTSs assembled *in vivo* from PCR fragments, we envision that RoCi will be particularly useful in high through-put discovery projects. This could be hunts for genes coding for novel enzymes or in mutational protein optimization schemes where many variants need to be produced and characterized. Since the growth volume in these types of experiments is low, the higher yields delivered by RoCi strains, as compared to single-copy strains, may significantly increase the chance of finding hits in a gene library simply because more genes, or gene variants, produce products in amounts that are sufficient for the products to be functionally characterized.

Most of our analyzes were performed in *A. nidulans* strains, but RoCi is not specific to this fungus as we obtained similar results with *A. niger* and *A. oryzae* as target organisms for RoCi. To this end we note that a recent CRISPR experiments using a c-GTS in an NHEJ proficient strain of *A. oryzae* produced transformants containing multi-copy integration events in the targeted locus (68). However, the orientation of the individual repeat structures in the gene clusters were not analyzed, and it is therefore not clear whether they were inserted via a RoCi event or not. Lastly, amplification of the 2µ circle of *S. cerevisiae* depends on double rolling-circle replication (69,70) and is therefore likely that RoCi will work in this important cell factory too. Certainly, it would not be surprising if RoCi also worked in non-fungal organisms. Moreover, we envision that RoCi can easily and advantageously be combined with other strategies to enhance gene expression, e.g. by employing strong synthetic promoters and terminators (71,72) and therefore serve as a popular addition to the growing synthetic biology toolbox for fungi.

## Acknowledgements

We thank Morten Kielland-Brandt and Jens Frisvad for the many insightful discussions and the Eukaryotic Molecular Cell Biology group for their support throughout the development of the project. This study was supported by the Innovation Fund Denmark (Grant Numbers 6150-00031B.9 and 0224-00121B).

## References

1. Liu D, Garrigues S, de Vries RP. Heterologous protein production in filamentous fungi. Appl Microbiol Biotechnol. 2023 Aug 5;107(16):5019–33.

2. Meyer V, Andersen MR, Brakhage AA, Braus GH, Caddick MX, Cairns TC, et al. Current challenges of research on filamentous fungi in relation to human welfare and a sustainable bio-economy: a white paper. Fungal Biol Biotechnol. 2016 Dec 31;3(1):6.

3. Hoffmeister D, Keller NP. Natural products of filamentous fungi: enzymes, genes, and their regulation. Nat Prod Rep. 2007 Apr;24(2):393–416.

4. Cairns TC, Nai C, Meyer V. How a fungus shapes biotechnology: 100 years of Aspergillus niger research. Fungal Biol Biotechnol. 2018;5:13.

5. Madhavan A, Arun KB, Sindhu R, Alphonsa Jose A, Pugazhendhi A, Binod P, et al. Engineering interventions in industrial filamentous fungal cell factories for biomass valorization. Bioresour Technol. 2022 Jan;344(Pt A):126209.

6. Nevalainen H, Peterson R. Making recombinant proteins in filamentous fungi-are we expecting too much? Front Microbiol. 2014;5:75.

7. Wang Q, Zhong C, Xiao H. Genetic Engineering of Filamentous Fungi for Efficient Protein Expression and Secretion. Front Bioeng Biotechnol. 2020 Apr 8;8.

8. Spiro RG. Glucose residues as key determinants in the biosynthesis and quality control of glycoproteins with N-linked oligosaccharides. J Biol Chem. 2000 Nov 17;275(46):35657–60.

9. Nevalainen KMH, Te’o VSJ, Bergquist PL. Heterologous protein expression in filamentous fungi. Trends Biotechnol. 2005 Sep;23(9):468–74.

10. Nødvig CS, Nielsen JB, Kogle ME, Mortensen UH. A CRISPR-Cas9 System for Genetic Engineering of Filamentous Fungi. PLoS One. 2015;10(7):e0133085.

11. Liu R, Chen L, Jiang Y, Zhou Z, Zou G. Efficient genome editing in filamentous fungus Trichoderma reesei using the CRISPR/Cas9 system. Cell Discov. 2015 May 12;1(1):15007.

12. Vanegas KG, Jarczynska ZD, Strucko T, Mortensen UH. Cpf1 enables fast and efficient genome editing in Aspergilli. Fungal Biol Biotechnol. 2019;6:6.

13. Katayama T, Tanaka Y, Okabe T, Nakamura H, Fujii W, Kitamoto K, et al. Development of a genome editing technique using the CRISPR/Cas9 system in the industrial filamentous fungus Aspergillus oryzae. Biotechnol Lett. 2016 Apr 19;38(4):637–42.

14. Leynaud-Kieffer LMC, Curran SC, Kim I, Magnuson JK, Gladden JM, Baker SE, et al. A new approach to Cas9-based genome editing in Aspergillus niger that is precise, efficient and selectable. PLoS One. 2019 Jan 17;14(1):e0210243.

15. Pohl C, Kiel JAKW, Driessen AJM, Bovenberg RAL, Nygård Y. CRISPR/Cas9 Based Genome Editing of Penicillium chrysogenum. ACS Synth Biol. 2016 Jul 15;5(7):754–64.

16. Nødvig CS, Hoof JB, Kogle ME, Jarczynska ZD, Lehmbeck J, Klitgaard DK, et al. Efficient oligo nucleotide mediated CRISPR-Cas9 gene editing in Aspergilli. Fungal Genet Biol. 2018 Jun;115:78–89.

17. Jarczynska ZD, Garcia Vanegas K, Deichmann M, Nørskov Jensen C, Scheeper MJ, Futyma ME, et al. A Versatile in Vivo DNA Assembly Toolbox for Fungal Strain Engineering. ACS Synth Biol. 2022 Oct 21;11(10):3251–63.

18. Verdoes JC, Punt PJ, Stouthamer AH, van den Hondel CA. The effect of multiple copies of the upstream region on expression of the Aspergillus niger glucoamylase-encoding gene. Gene. 1994 Aug 5;145(2):179–87.

19. Jeenes DJ, Mackenzie DA, Archer DB. Transcriptional and post-transcriptional events affect the production of secreted hen egg white lysozyme byAspergillus niger. Transgenic Res. 1994 Sep;3(5):297–303.

20. Wang G, Huang M, Nielsen J. Exploring the potential of Saccharomyces cerevisiae for biopharmaceutical protein production. Curr Opin Biotechnol. 2017 Dec;48:77–84.

21. Baeshen NA, Baeshen MN, Sheikh A, Bora RS, Ahmed MMM, Ramadan HAI, et al. Cell factories for insulin production. Microb Cell Fact. 2014 Oct 2;13:141.

22. Kjeldsen T, Andersen AS, Hubálek F, Johansson E, Kreiner FF, Schluckebier G, et al. Molecular engineering of insulin for recombinant expression in yeast. Trends Biotechnol. 2024 Apr;42(4):464–78.

23. Aleksenko A, Clutterbuck AJ. Autonomous plasmid replication in Aspergillus nidulans: AMA1 and MATE elements. Fungal Genet Biol. 1997 Jun;21(3):373–87.

24. Gems D, Johnstone IL, Clutterbuck AJ. An autonomously replicating plasmid transforms Aspergillus nidulans at high frequency. Gene. 1991 Feb;98(1):61–7.

25. Hansen BG, Salomonsen B, Nielsen MT, Nielsen JB, Hansen NB, Nielsen KF, et al. Versatile enzyme expression and characterization system for Aspergillus nidulans, with the Penicillium brevicompactum polyketide synthase gene from the mycophenolic acid gene cluster as a test case. Appl Environ Microbiol. 2011 May;77(9):3044–51.

26. Mikkelsen MD, Buron LD, Salomonsen B, Olsen CE, Hansen BG, Mortensen UH, et al. Microbial production of indolylglucosinolate through engineering of a multi-gene pathway in a versatile yeast expression platform. Metab Eng. 2012 Mar;14(2):104–11.

27. Dujon B. The yeast genome project: what did we learn? Trends Genet. 1996 Jul;12(7):263–70.

28. Lopes TS, Hakkaart GJ, Koerts BL, Raué HA, Planta RJ. Mechanism of high-copy-number integration of pMIRY-type vectors into the ribosomal DNA of Saccharomyces cerevisiae. Gene. 1991 Aug 30;105(1):83–90.

29. Semkiv M V., Dmytruk K V., Sibirny AA. Development of a system for multicopy gene integration in Saccharomyces cerevisiae. J Microbiol Methods. 2016 Jan;120:44–9.

30. Lee FWF, Silva NA Da. Improved efficiency and stability of multiple cloned gene insertions at the δ sequences of Saccharomyces cerevisiae. Appl Microbiol Biotechnol. 1997 Sep 26;48(3):339–45.

31. Wang X, Wang Z, Da Silva NA. G418 Selection and stability of cloned genes integrated at chromosomal delta sequences of Saccharomyces cerevisiae. Biotechnol Bioeng. 1996 Jan 5;49(1):45–51.

32. Wang Z, Da Silva NA. Improved protein synthesis and secretion through medium enrichment in a stable recombinant yeast strain. Biotechnol Bioeng. 1993 Jun 5;42(1):95–102.

33. Lopes TS, de Wijs IJ, Steenhauer SI, Verbakel J, Planta RJ. Factors affecting the mitotic stability of high-copy-number integration into the ribosomal DNA of Saccharomyces cerevisiae. Yeast. 1996 Apr;12(5):467–77.

34. Yelton MM, Hamer JE, Timberlake WE. Transformation of Aspergillus nidulans by using a trpC plasmid. Proc Natl Acad Sci U S A. 1984 Mar;81(5):1470–4.

35. Ballance DJ, Turner G. Development of a high-frequency transforming vector for Aspergillus nidulans. Gene. 1985;36(3):321–31.

36. Buxton FP, Gwynne DI, Davies RW. Transformation of Aspergillus niger using the argB gene of Aspergillus nidulans. Gene. 1985;37(1–3):207–14.

37. Kelly JM, Hynes MJ. Transformation of Aspergillus niger by the amdS gene of Aspergillus nidulans. EMBO J. 1985 Feb;4(2):475–9.

38. Wernars K, Goosen T, Wennekes LM, Visser J, Bos CJ, van den Broek HW, et al. Gene amplification in Aspergillus nidulans by transformation with vectors containing the amdS gene. Curr Genet. 1985 May;9(5):361–8.

39. Strucko T, Buron LD, Jarczynska ZD, Nødvig CS, Mølgaard L, Halkier BA, et al. CASCADE, a platform for controlled gene amplification for high, tunable and selection-free gene expression in yeast. Sci Rep. 2017 Jan 30;7:41431.

40. Arentshorst M, Kooloth Valappil P, Mózsik L, Regensburg-Tuïnk TJG, Seekles SJ, Tjallinks G, et al. A CRISPR/Cas9-based multicopy integration system for protein production in Aspergillus niger. FEBS J. 2023 Nov 27;290(21):5127–40.

41. Kaminskyj SGW. Fundamentals of growth, storage, genetics and microscopy of Aspergillus nidulans. Fungal Genet Rep. 2001 Sep 1;48(1):25–31.

42. Cove DJ. The induction and repression of nitrate reductase in the fungus Aspergillus nidulans. Biochim Biophys Acta. 1966 Jan;113(1):51–6.

43. Nørholm MHH. A mutant Pfu DNA polymerase designed for advanced uracil-excision DNA engineering. BMC Biotechnol. 2010 Mar 16;10:21.

44. Nour-Eldin HH, Geu-Flores F, Halkier BA. USER Cloning and USER Fusion: The Ideal Cloning Techniques for Small and Big Laboratories. In 2010. p. 185–200.

45. Jarczynska ZD, Rendsvig JKH, Pagels N, Viana VR, Nødvig CS, Kirchner FH, et al. DIVERSIFY: A Fungal Multispecies Gene Expression Platform. ACS Synth Biol. 2021 Mar 19;10(3):579–88.

46. Nielsen ML, Albertsen L, Lettier G, Nielsen JB, Mortensen UH. Efficient PCR-based gene targeting with a recyclable marker for Aspergillus nidulans. Fungal Genet Biol. 2006 Jan;43(1):54–64.

47. Gautam A. Phenol-Chloroform DNA Isolation Method. In 2022. p. 33–9.

48. Green MR, Sambrook J. Precipitation of DNA with Isopropanol. Cold Spring Harb Protoc. 2017 Aug 1;2017(8):pdb.prot093385.

49. Church GM, Gilbert W. Genomic sequencing. Proc Natl Acad Sci U S A. 1984 Apr;81(7):1991–5.

50. Schoustra SE, Debets AJM, Slakhorst M, Hoekstra RF. Reducing the cost of resistance; experimental evolution in the filamentous fungus Aspergillus nidulans. J Evol Biol. 2006 Jul;19(4):1115–27.

51. Smedsgaard J. Micro-scale extraction procedure for standardized screening of fungal metabolite production in cultures. J Chromatogr A. 1997 Jan;760(2):264–70.

52. Jefferson RA. Assaying chimeric genes in plants: The GUS gene fusion system. Plant Mol Biol Report. 1987 Dec;5(4):387–405.

53. Bradford MM. A rapid and sensitive method for the quantitation of microgram quantities of protein utilizing the principle of protein-dye binding. Anal Biochem. 1976 May 7;72:248–54.

54. Ait Saada A, Guo W, Costa AB, Yang J, Wang J, Lobachev KS. Widely spaced and divergent inverted repeats become a potent source of chromosomal rearrangements in long single-stranded DNA regions. Nucleic Acids Res. 2023 May 8;51(8):3722–34.

55. Lai PJ, Lim CT, Le HP, Katayama T, Leach DRF, Furukohri A, et al. Long inverted repeat transiently stalls DNA replication by forming hairpin structures on both leading and lagging strands. Genes to Cells. 2016 Feb 6;21(2):136–45.

56. Voineagu I, Narayanan V, Lobachev KS, Mirkin SM. Replication stalling at unstable inverted repeats: interplay between DNA hairpins and fork stabilizing proteins. Proc Natl Acad Sci U S A. 2008 Jul 22;105(29):9936–41.

57. Fiddy C, Trinci APJ. Mitosis, Septation, Branching and the Duplication Cycle in Aspergillus nidulans. J Gen Microbiol. 1976 Dec 1;97(2):169–84.

58. Harris SD. The Duplication Cycle inAspergillus nidulans. Fungal Genetics and Biology. 1997 Aug;22(1):1–12.

59. Momany M. Polarity in filamentous fungi: establishment, maintenance and new axes. Curr Opin Microbiol. 2002 Dec;5(6):580–5.

60. Erhart E, Hollenberg CP. The presence of a defective LEU2 gene on 2 mu DNA recombinant plasmids of Saccharomyces cerevisiae is responsible for curing and high copy number. J Bacteriol. 1983 Nov;156(2):625–35.

61. Wong YY, Moon A, Duffin R, Barthet-Barateig A, Meijer HA, Clemens MJ, et al. Cordycepin Inhibits Protein Synthesis and Cell Adhesion through Effects on Signal Transduction. Journal of Biological Chemistry. 2010 Jan;285(4):2610–21.

62. Liao Y, Ling J, Zhang G, Liu F, Tao S, Han Z, et al. Cordycepin induces cell cycle arrest and apoptosis by inducing DNA damage and up-regulation of p53 in Leukemia cells. Cell Cycle. 2015 Mar 4;14(5):761–71.

63. Rose KM, Bell LE, Jacob ST. Specific inhibition of chromatin-associated poly(A) synthesis in vitro by cordycepin 5′-triphosphate. Nature. 1977 May;267(5607):178–80.

64. Kaczka EA, Dulaney EL, Gitterman CO, Boyd Woodruff H, Folkers K. Isolation and inhibitory effects on KB cell cultures of 3′-deoxyadenosine from Aspergillus nidulans (Eidam) wint. Biochem Biophys Res Commun. 1964 Jan;14(5):452–5.

65. Xia Y, Luo F, Shang Y, Chen P, Lu Y, Wang C. Fungal Cordycepin Biosynthesis Is Coupled with the Production of the Safeguard Molecule Pentostatin. Cell Chem Biol. 2017 Dec;24(12):1479–1489.e4.

66. Masuda M, Urabe E, Sakurai A, Sakakibara M. Production of cordycepin by surface culture using the medicinal mushroom Cordyceps militaris. Enzyme Microb Technol. 2006 Aug;39(4):641–6.

67. Gaillard H, García-Muse T, Aguilera A. Replication stress and cancer. Nat Rev Cancer. 2015 May 24;15(5):276–89.

68. Katayama T, Nakamura H, Zhang Y, Pascal A, Fujii W, Maruyama JI. Forced Recycling of an AMA1-Based Genome-Editing Plasmid Allows for Efficient Multiple Gene Deletion/Integration in the Industrial Filamentous Fungus Aspergillus oryzae. Appl Environ Microbiol. 2019 Feb 1;85(3).

69. Volkert FC, Broach JR. Site-specific recombination promotes plasmid amplification in yeast. Cell. 1986 Aug;46(4):541–50.

70. Futcher AB. Copy number amplification of the 2 μm circle plasmid of Saccharomyces cerevisiae. J Theor Biol. 1986 Mar;119(2):197–204.

71. Deng J, Wu Y, Zheng Z, Chen N, Luo X, Tang H, et al. A synthetic promoter system for well-controlled protein expression with different carbon sources in Saccharomyces cerevisiae. Microb Cell Fact. 2021 Dec 18;20(1):202.

72. Curran KA, Morse NJ, Markham KA, Wagman AM, Gupta A, Alper HS. Short Synthetic Terminators for Improved Heterologous Gene Expression in Yeast. ACS Synth Biol. 2015 Jul 17;4(7):824–32.

