## Supplementary material for "RoCi - A Single Step Multi-Copy Integration System Based on Rolling-Circle Replication": Supplementary RoCi.pdf

**The supplementary information contains 9 Figures and 6 Tables:**

**Figures**


- **Figure S1:** Bioreactor cultivation conditions.
- **Figure S2:** Visible- and red fluorescent light images of RoCi strains.
- **Figure S3:** RoCi complex repair schemes.
- **Figure S4:** Phenotype verification from RoCi experiment with pDIV088 & pDIV089**.**
- **Figure S5:** VILBER-FX images and 3D fluorescence spectra from pre-formed c-GTS versus c-GTS made by *in vivo* assembly comparison experiments.
- **Figure S6:** Solid media stability experiments with mRFP production.
- **Figure S7:** Single-copy versus multi-copy *pyrG-d::mRFP* strains. Growth and fluorescence assessment.
- **Figure S8:** Plug extraction plates and standard curve for cordycepin titter calculation.
- **Figure S9:** Illustration of *uidA*-GEC/*mRFP* vector assembly *in vivo.*


**Tables**


- **Table S1:** Plasmids used in this study.
- **Table S2:** List of fungal strains.
- **Table S3:** Primers used in this study.
- **Table S4:** Protospacers used in this study.
- **Table S5:** Probes for ddPCR used in this study.
- **Table S6:** Copy-number data from pre-formed c-GTS versus c-GTS made by *in vivo* assembly comparison experiments.

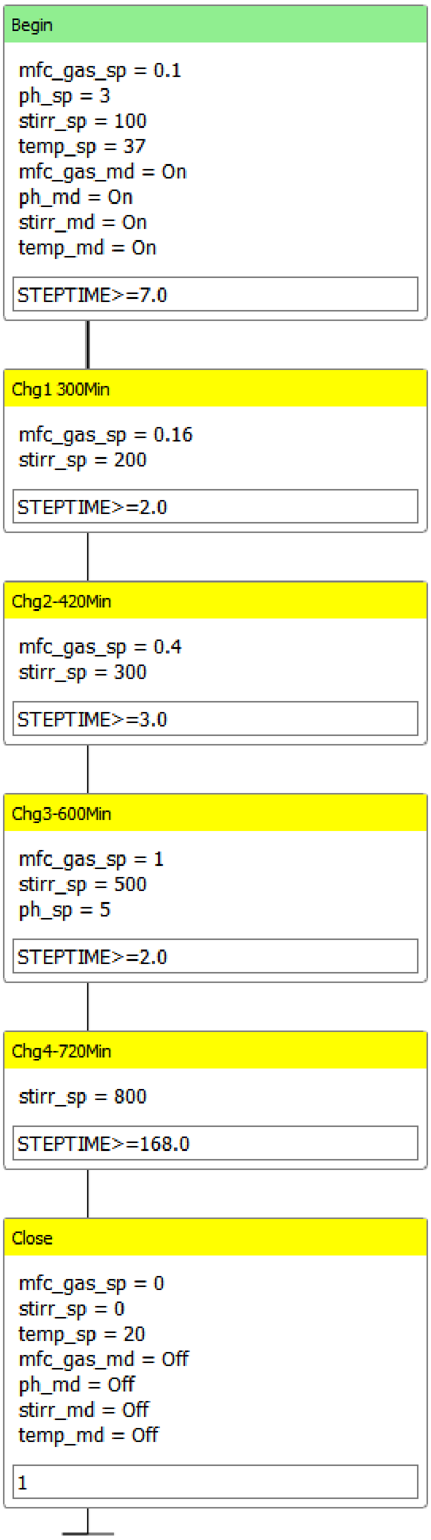


**Figure S1. Bioreactor cultivation conditions.** Temperature (temp_sp), pH (ph_sp), aeration (mfc_gas_sp) and stirring (stirr_sp) conditions of each cultivation stage are shown. In step Chg4-720Min the STEPTIME was set for 168 hour, but the cultivation was stopped at 38 hour (total time).

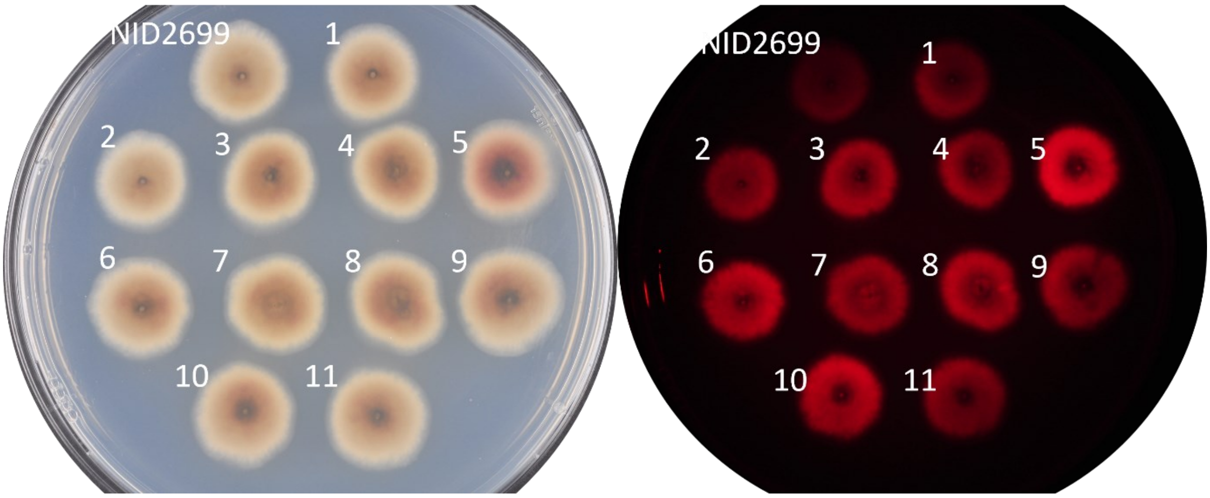


**Figure S2. RoCi strains analyzed by ddPCR and Southern-blot**. Visible- and red fluorescence light images of *A. nidulans* RoCi strains, stabbed in a 14.5 cm Petri dish.

A

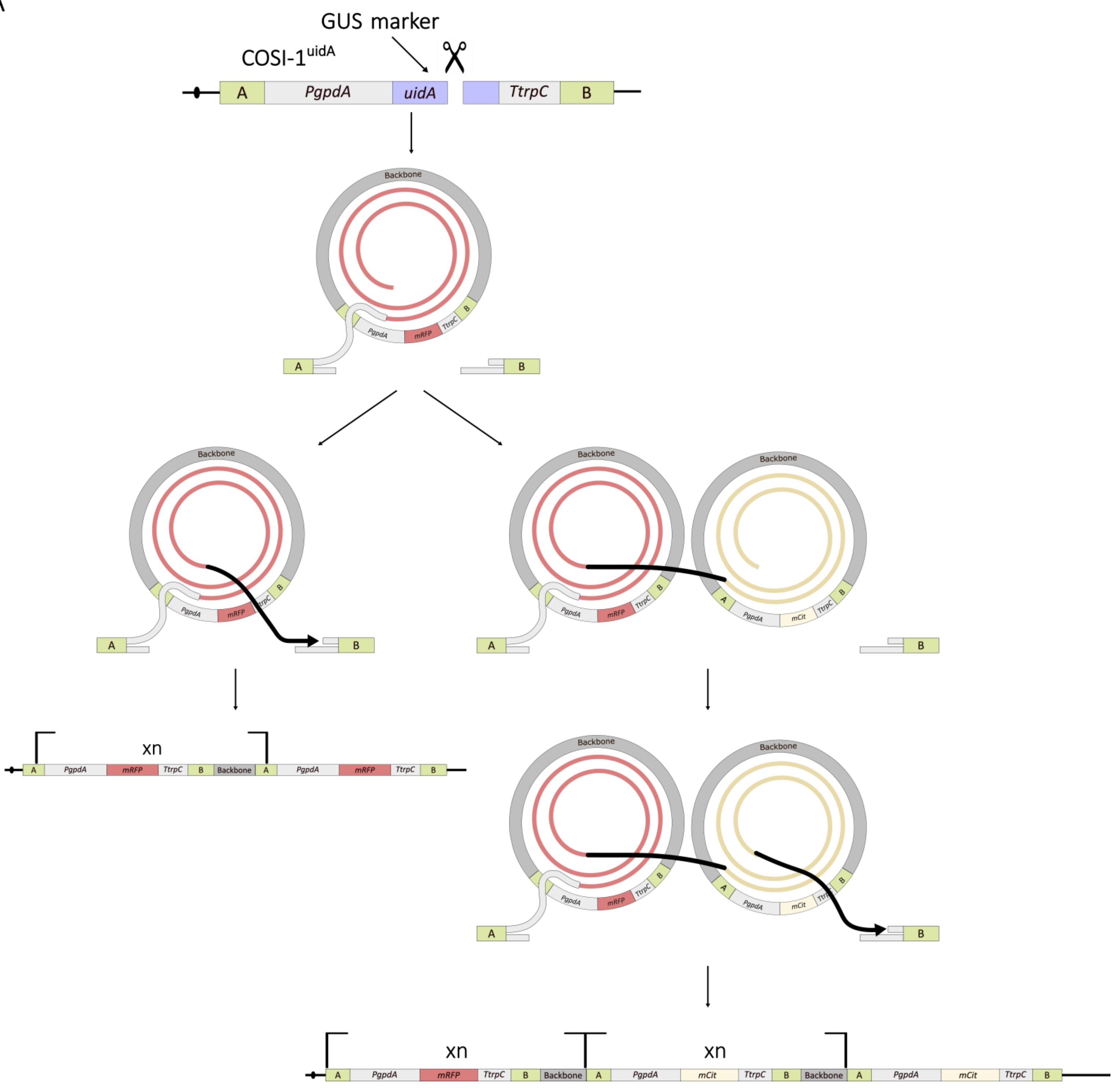

GUS marker
COSI-1uidA
A
PgpdA
uidA
TtrpC
B
Backbone
PgpdA
mRFP
TtrpC
mCit
xn

B

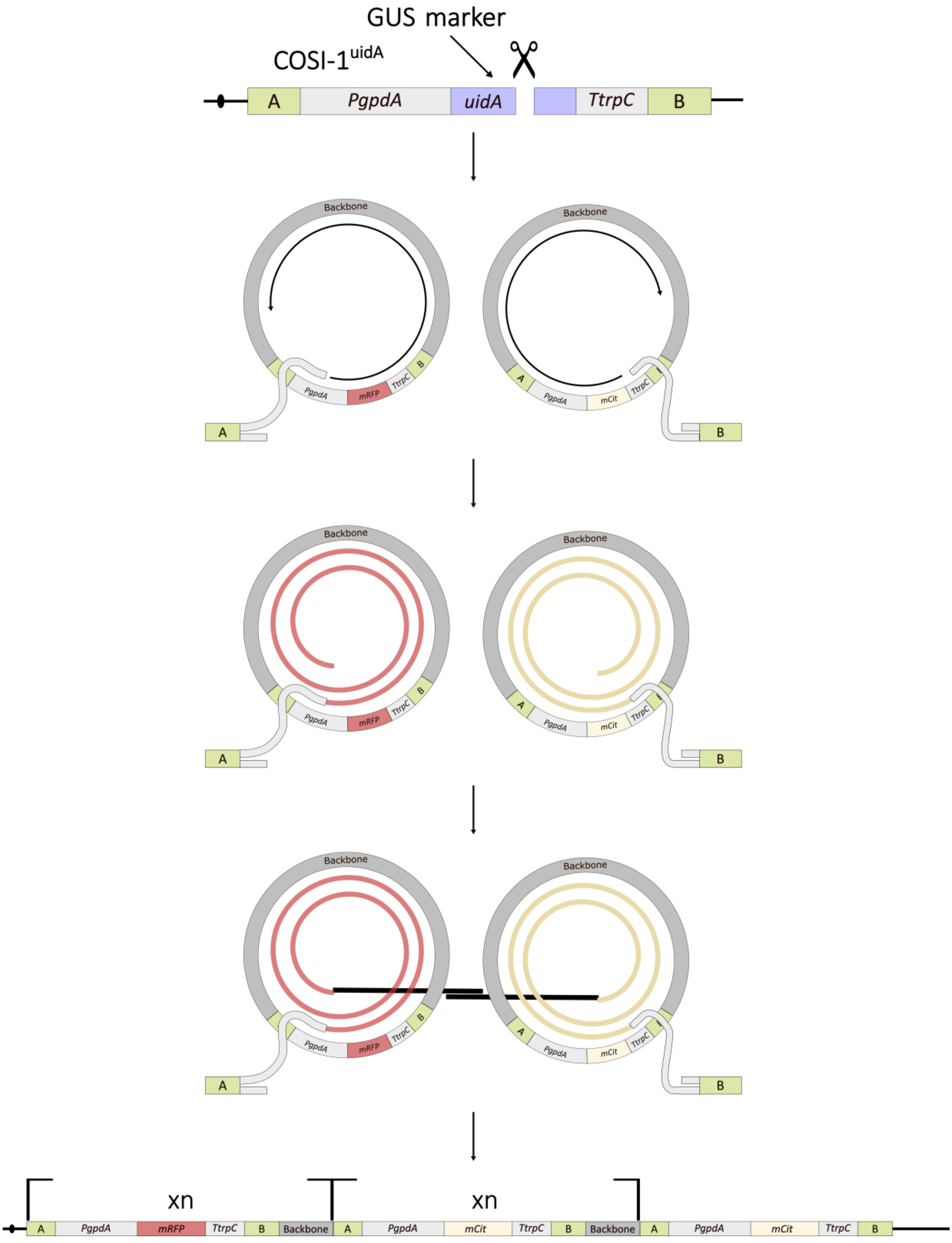

GUS marker
COSI-1uidA
A
PgpdA
uidA
TtrpC
B
Backbone
PgpdA
mRFP
TtrpC
mCit
xn
Backbone

**Figure S3. Complex RoCi integration events involving more than one strand invasion:** (**A**) After invading the c-GTS, the elongating strand has two options when the replication fork collapses, either to do the second end-capture directly (left), which will result in the integration of only one type of GEC (*mRFP* in this case); or invade another c-GTS for further extension and eventually do the second end-capture, which could potentially result in the integration of two different GECs. (**B**) The two liberated 3' ends can invade two different c-GTS, possibly integrating two different GECs, and after extension they can fuse by annealing to repair the DSB. As a result of this event either one or both GECs could be integrated.

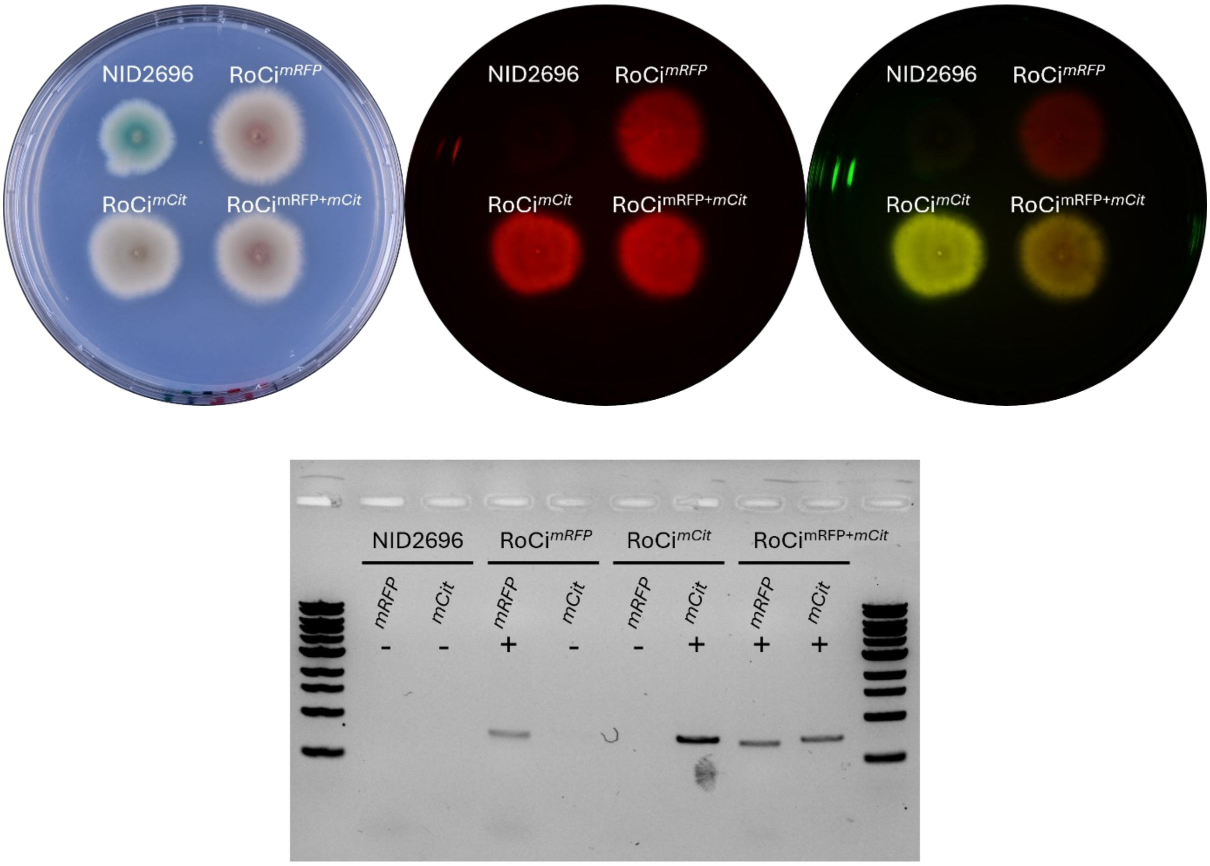


**Figure S4. Phenotype verification from RoCi experiment with pDIV088 & pDIV089.** The three different populations resulting from the transformation of NID2696 with plasmids bearing a *mCit* GEC (pDIV088) and a *mRFP* GEC (pDIV089) were stabbed in a MM + X-gluc plate and imaged under visible-, red fluorescent-, and yellow fluorescent light. Under red fluorescent light, all colonies appear red. However, under yellow fluorescent light, the strains bearing only *mRFP* copies appear red (RoCi$^{mRFP}$), the strains bearing only *mCit* copies appear yellow (RoCi$^{mCit}$) and strains bearing both GECs appear orange (RoCi $^{\mathrm{mRFP+mCit}}$). The phenotype scoring was confirmed by PCR looking for the presence of the corresponding genes.

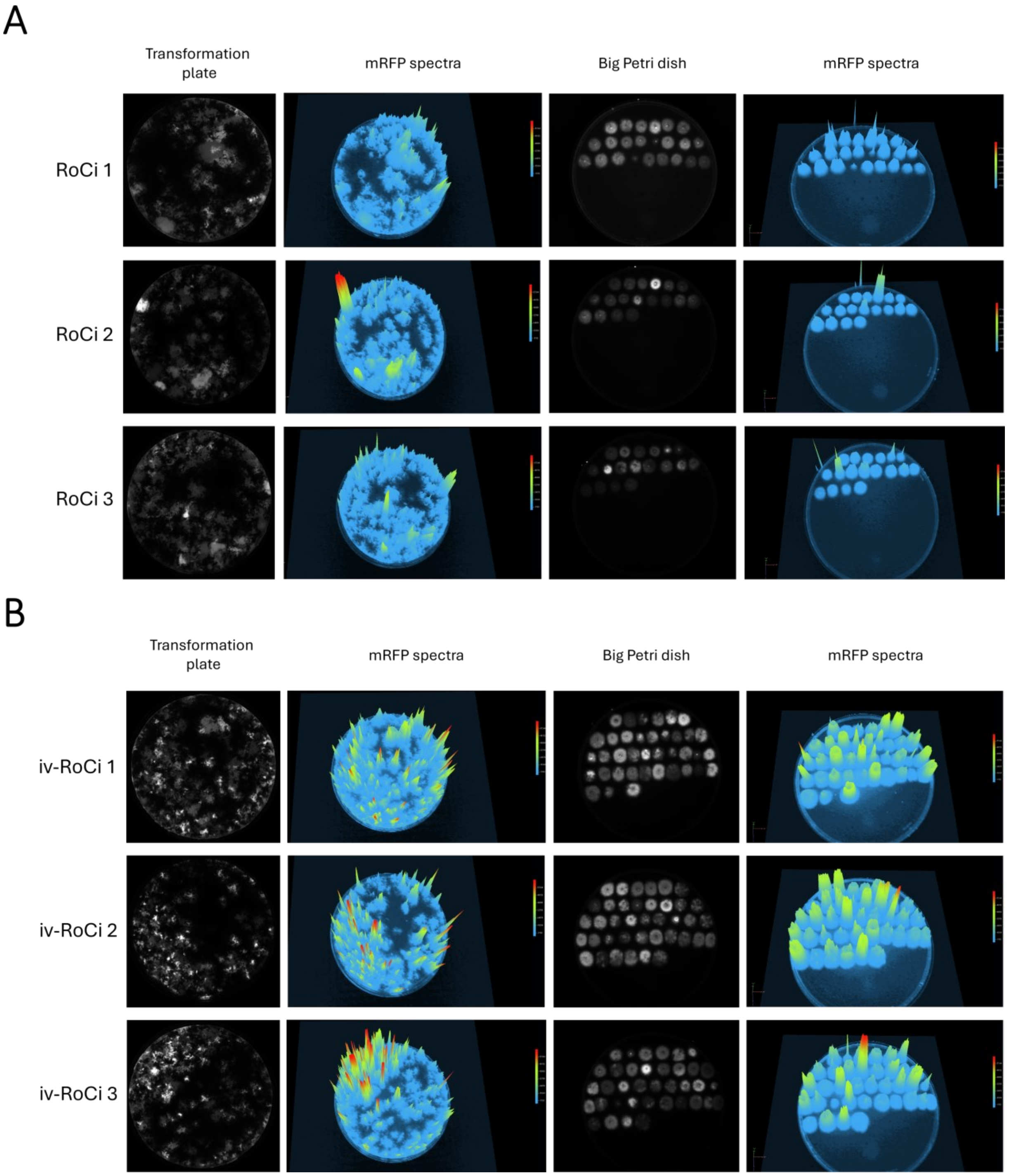


**Figure S5. Pre-formed c-GTS versus c-GTS made by *in vivo* assembly.** Transformation plates and fluorescent isolates were imaged under red fluorescence light setup using the Vilber Fusion FX (Exposure time 1.3 s). Finally, a 3D fluorescent spectra of each plate was made to compare fluorescence intensities. Images from panel **A** correspond to the transformations done with a pre-formed c-GTS (RoCi) and images in panel **B** correspond to the transformations done with c-GTS made by in vivo assembly (iv-RoCi).

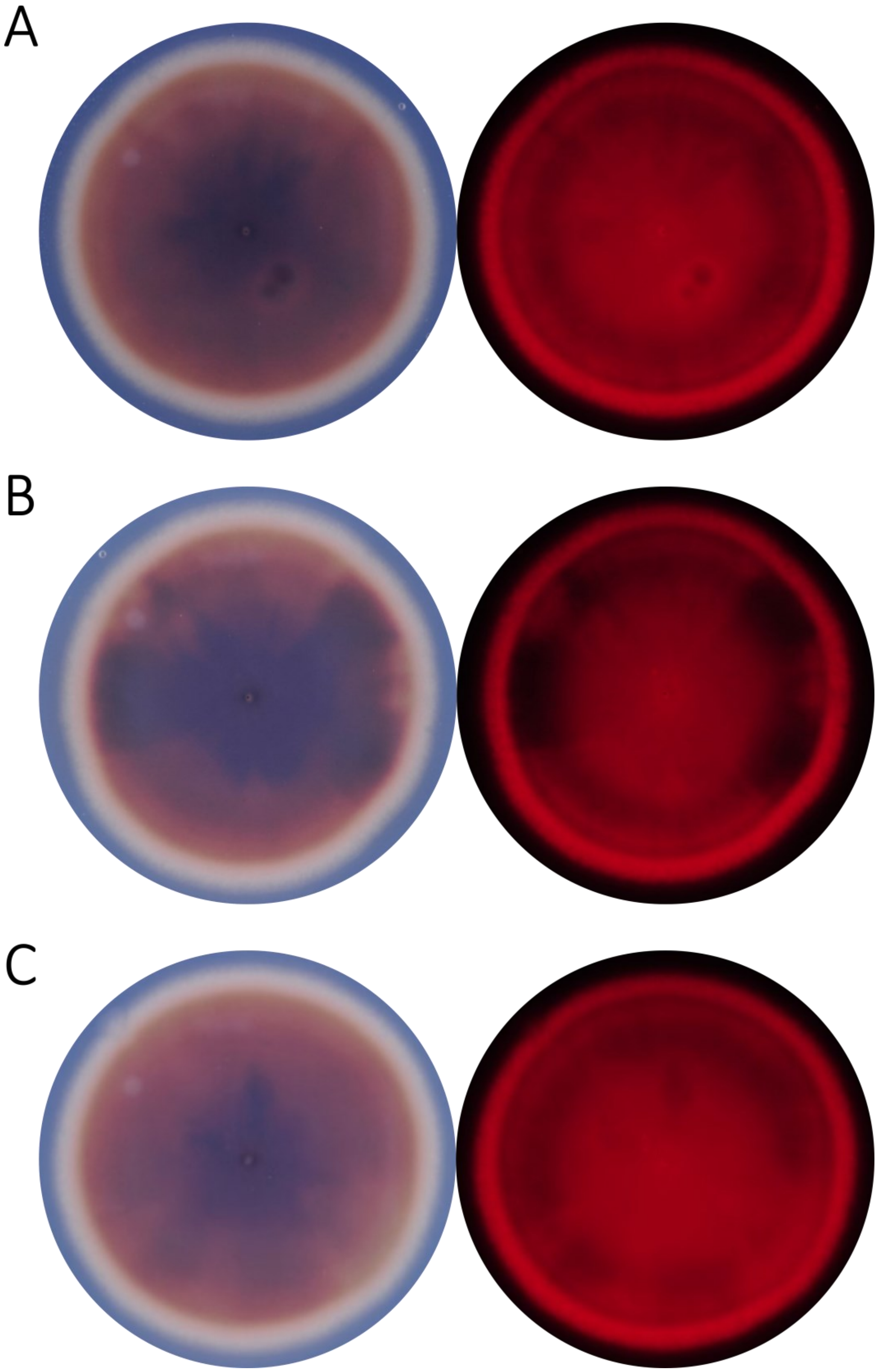


**Figure S6. Solid media stability test.** The strain sDIV0526 (23 *mRFP* GEC copies) was inoculated in 14.5 cm Petri dishes in triplicate (A, B & C).

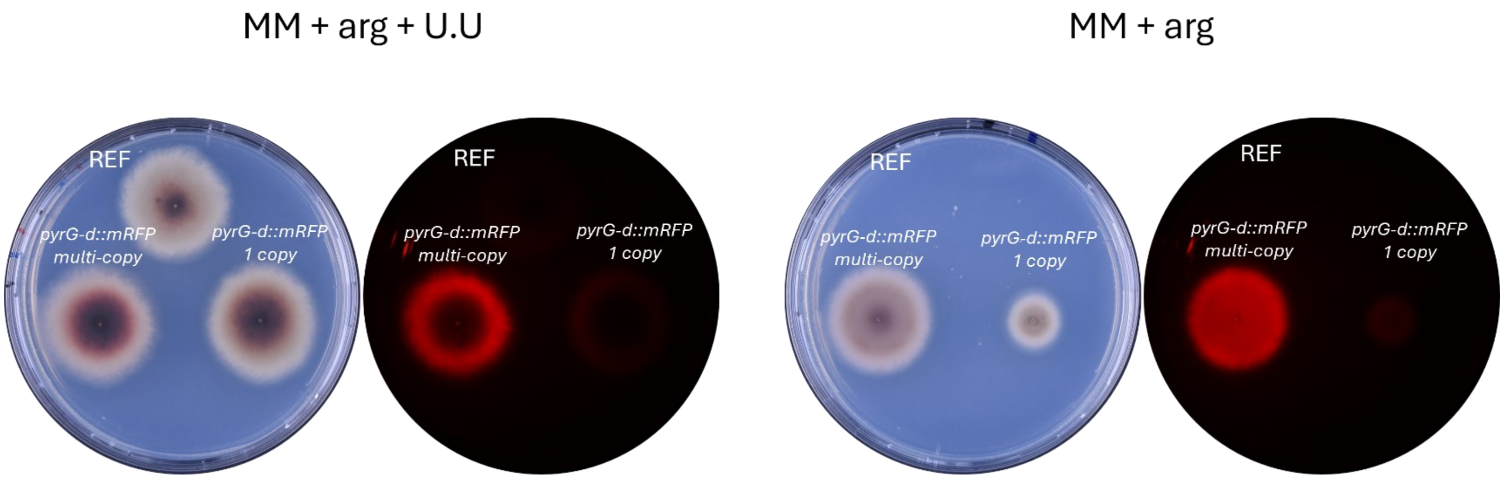


**Figure S7. Single-copy versus multi-copy *pyrG-d::mRFP* strains**. Growth comparison of the REF strain NID2696, single copy *pyrG-d::RFP* GEC strain and the multi-copy *pyrG-d::RFP* GEC strain with and without uridine and uracil (U.U) supplementation. Plates were imaged in visible- and red fluorescent light.

A

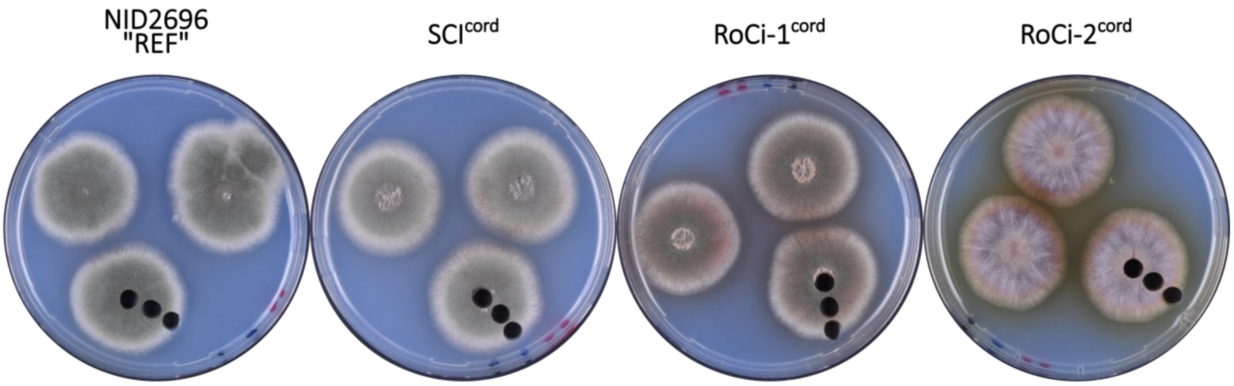


B

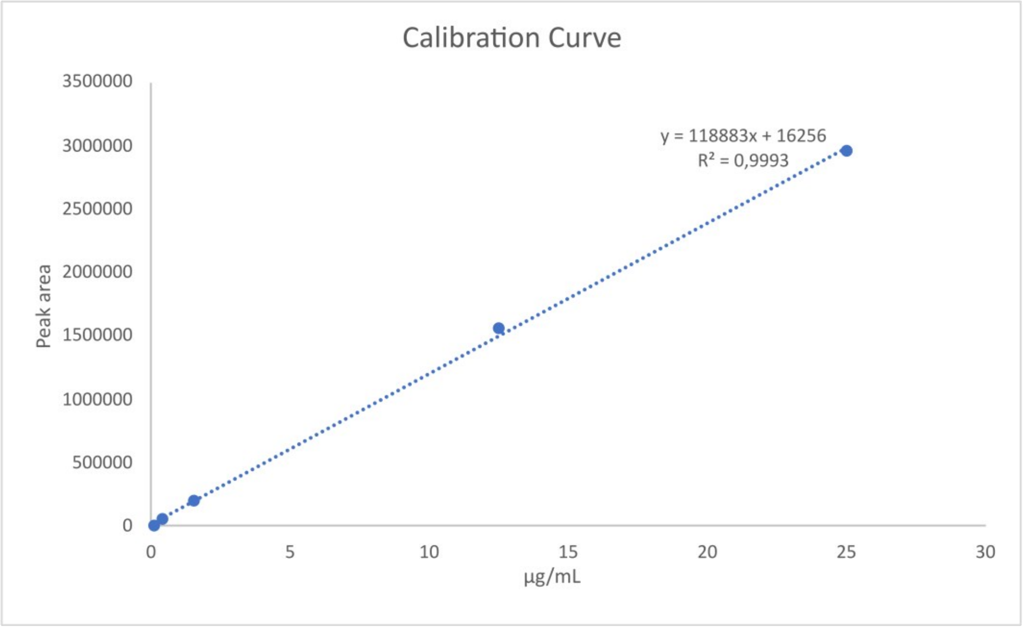


**Figure S8. Plug extraction plates and standard curve for cordycepin titter calculation.** (**A**) Plates from the plug extractions for relative quantification of cordycepin. (**B**) Calibration curve to calculate cordycepin concentration in plug extraction samples.

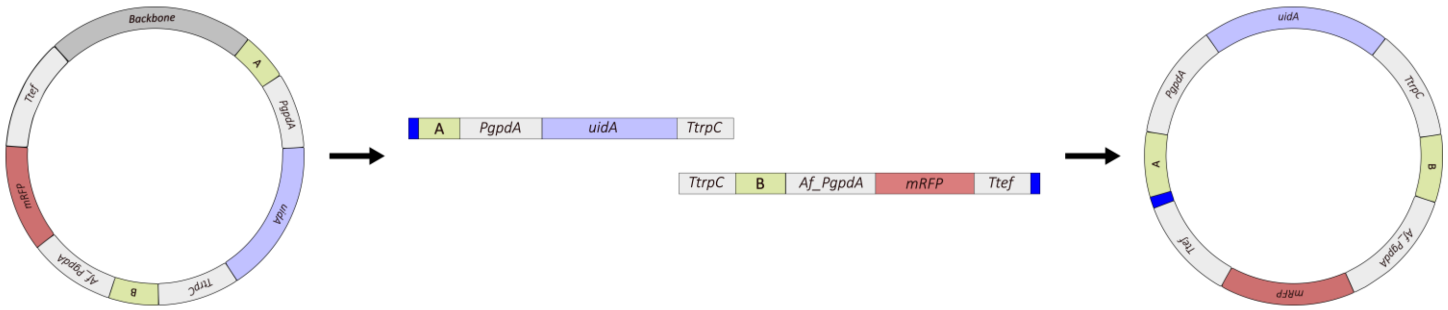


**Figure S9. Circular *uidA*-GEC/*mRFP* vector assembly by *in vivo* recombination**. Parting from plasmid pDIV0941 as PCR template, two overlapping PCR fragments are generated covering the entire *TtrpC* sequence and the "A" and "B" ends of the substrates are extended with 50 bp assembly tags (dark blue) by primer overhangs. The two fragments have two overlapping sequences that allow for c-GTS assembly *in vivo*.

**Table S1:** Plasmids used in this study.

| Plasmid ID | Description[1] | Reference |
|---|---|---|
| pU2002A | PacI/Nt.BbvCI | Hansen et al., (1) |
| pFC330 | pAMA1-pyrG-Cas9-PacI/Nt.BbvCI | Nødvig et al., (2) |
| pDIV068 | A-*An*_PgpdA-PacI-An-T*trpC*-B | Jarczynska et al., (3) |
| pDIV073 | pFC330::*uidA*-gRNA2 | Jarczynska et al., (3) |
| pDIV089 | A-*An*_P*gpdA-mRFP-An*_T*trpC-B* | Jarczynska et al., (3) |
| pDIV131 | A-PacI-B | Jarczynska et al., (3) |
| pDIV088 | A-*An*_P*gpdA-mCitrine-An*-T*trpC*-B | This study |
| pDIV1050 | A-*An*_P*tef*-$cns_1$(EAAAK)$_4$$cns_2$-*An*_T*tef*-B | This study |
| pDIV1051 | A-*Af_pyrGd-An*_P*gpdA-mRFP-An*_T*trpC*-B | This study |
| pDIV1049 | pFC330::*mCit*-gRNA1 | This study |
| pDIV0941 | A-*An_PgpdA-uidA-An_TrpC*-B-*Af_PgpdA(0.8 kb)-mRFP-An_Ttef* | This study |

[1]An- *Aspergillus nidulans*; Af- *Aspergillus fumigatus*

**Table S2:** List of fungal strains.

| **Strain ID** | **"Alias"**[1] | **Description**[1] | **Reference** |
|---|---|---|---|
| NID2696 | COSI-1$^{uidA}$ | *argB2; pyrG89; veA1; nkuAΔ; IS1::*A-*An*-P*gpdA-uidA-An*_T*trpC*-B | Jarczynska et al., (3) |
| NID2699 | SCI | *argB2; pyrG89; veA1; nkuAΔ; IS1::*A-*An*-P*gpdA-mRFP-An*_T*trpC*-B | Jarczynska et al., (3) |
| sDIV0520 | RoCi-1$^{mRFP}$ | *argB2; pyrG89; veA1; nkuAΔ; IS1::(*A-*An*-P*gpdA-mRFP-An_TtrpC-B-Backbone*)$_2$ | This study |
| | RoCi-2 $^{mRFP}$ | *argB2; pyrG89; veA1; nkuAΔ; IS1::(*A-*An*-P*gpdA-mRFP-An_TtrpC-B-Backbone*)$_2$ | This study |
| | RoCi-3 $^{mRFP}$ | *argB2; pyrG89; veA1; nkuAΔ; IS1::(*A-*An*-P*gpdA-mRFP-An_TtrpC-B-Backbone*)$_2$ | This study |
| | RoCi-4 $^{mRFP}$ | *argB2; pyrG89; veA1; nkuAΔ; IS1::(*A-*An*-P*gpdA-mRFP-An_TtrpC-B-Backbone*)$_2$ | This study |
| sDIV0521 | RoCi-5 $^{mRFP}$ | *argB2; pyrG89; veA1; nkuAΔ; IS1::(*A-*An*-P*gpdA-mRFP-An_TtrpC-B-Backbone*)$_{11}$ | This study |
| sDIV0522 | RoCi-6 $^{mRFP}$ | *argB2; pyrG89; veA1; nkuAΔ; IS1::(*A-*An*-P*gpdA-mRFP-An_TtrpC-B-Backbone*)$_3$ | This study |
| | RoCi-7 $^{mRFP}$ | *argB2; pyrG89; veA1; nkuAΔ; IS1::(*A-*An*-P*gpdA-mRFP-An_TtrpC-B-Backbone*)$_2$ | This study |
| | RoCi-8 $^{mRFP}$ | *argB2; pyrG89; veA1; nkuAΔ; IS1::(*A-*An*-P*gpdA-mRFP-An_TtrpC-B-Backbone*)$_3$ | This study |
| sDIV0523 | RoCi-9 $^{mRFP}$ | *argB2; pyrG89; veA1; nkuAΔ; IS1::(*A-*An*-P*gpdA-mRFP-An_TtrpC-B-Backbone*)$_4$ | This study |
| sDIV0524 | RoCi-10 $^{mRFP}$ | *argB2; pyrG89; veA1; nkuAΔ; IS1::(*A-*An*-P*gpdA-mRFP-An_TtrpC-B-Backbone*)$_5$ | This study |
| | RoCi-11 $^{mRFP}$ | *argB2; pyrG89; veA1; nkuAΔ; IS1::(*A-*An*-P*gpdA-mRFP-An_TtrpC-B-Backbone*)$_3$ | This study |
| | RoCi-1.1$^{mRFP}$ | *argB2; pyrG89; veA1; nkuAΔ; IS1::(*A-*An*-P*gpdA-mRFP-An_TtrpC-B-BAckbone)* $_n$ | This study |
| | RoCi -1.2$^{mRFP}$ | *argB2; pyrG89; veA1; nkuAΔ; IS1::(*A-*An*-P*gpdA-mRFP-An_TtrpC-B-BAckbone)* $_n$ | This study |

| | | | |
|---|---|---|---|
| | RoCi -1.3$^{mRFP}$ | *argB2; pyrG89; veA1; nkuAΔ; IS1::(*A-*An*-P*gpdA-mRFP-An_TtrpC-B-BAckbone)* $_{n}$ | This study |
| | RoCi -1.4$^{mRFP}$ | *argB2; pyrG89; veA1; nkuAΔ; IS1::(*A-*An*-P*gpdA-mRFP-An_TtrpC-B-BAckbone)* $_{n}$ | This study |
| | RoCi -1.5$^{mRFP}$ | *argB2; pyrG89; veA1; nkuAΔ; IS1::(*A-*An*-P*gpdA-mRFP-An_TtrpC-B-BAckbone)* $_{n}$ | This study |
| | RoCi -2.1$^{mRFP}$ | *argB2; pyrG89; veA1; nkuAΔ; IS1::(*A-*An*-P*gpdA-mRFP-An_TtrpC-B-BAckbone)* $_{n}$ | This study |
| | RoCi -2.2$^{mRFP}$ | *argB2; pyrG89; veA1; nkuAΔ; IS1::(*A-*An*-P*gpdA-mRFP-An_TtrpC-B-BAckbone)* $_{n}$ | This study |
| | RoCi -2.3$^{mRFP}$ | *argB2; pyrG89; veA1; nkuAΔ; IS1::(*A-*An*-P*gpdA-mRFP-An_TtrpC-B-BAckbone)* $_{n}$ | This study |
| | RoCi -2.4$^{mRFP}$ | *argB2; pyrG89; veA1; nkuAΔ; IS1::(*A-*An*-P*gpdA-mRFP-An_TtrpC-B-BAckbone)* $_{n}$ | This study |
| | RoCi -2.5$^{mRFP}$ | *argB2; pyrG89; veA1; nkuAΔ; IS1::(*A-*An*-P*gpdA-mRFP-An_TtrpC-B-BAckbone)* $_{n}$ | This study |
| | RoCi -3.1$^{mRFP}$ | *argB2; pyrG89; veA1; nkuAΔ; IS1::(*A-*An*-P*gpdA-mRFP-An_TtrpC-B-BAckbone)* $_{n}$ | This study |
| | RoCi -3.2$^{mRFP}$ | *argB2; pyrG89; veA1; nkuAΔ; IS1::(*A-*An*-P*gpdA-mRFP-An_TtrpC-B-BAckbone)* $_{n}$ | This study |
| | RoCi -3.3$^{mRFP}$ | *argB2; pyrG89; veA1; nkuAΔ; IS1::(*A-*An*-P*gpdA-mRFP-An_TtrpC-B-BAckbone)* $_{n}$ | This study |
| | RoCi -3.4$^{mRFP}$ | *argB2; pyrG89; veA1; nkuAΔ; IS1::(*A-*An*-P*gpdA-mRFP-An_TtrpC-B-BAckbone)* $_{n}$ | This study |
| | RoCi -3.5$^{mRFP}$ | *argB2; pyrG89; veA1; nkuAΔ; IS1::(*A-*An*-P*gpdA-mRFP-An_TtrpC-B-BAckbone)* $_{n}$ | This study |
| | iv-RoCi-1.1$^{mRFP}$ | *argB2; pyrG89; veA1; nkuAΔ; IS1::(*A-*An*-P*gpdA-mRFP-An_TtrpC-B*) $_{n}$ | This study |
| | iv-RoCi-1.2$^{mRFP}$ | *argB2; pyrG89; veA1; nkuAΔ; IS1::(*A-*An*-P*gpdA-mRFP-An_TtrpC-B*) $_{n}$ | This study |
| | iv-RoCi-1.3$^{mRFP}$ | *argB2; pyrG89; veA1; nkuAΔ; IS1::(*A-*An*-P*gpdA-mRFP-An_TtrpC-B*) $_{n}$ | This study |

| | | | |
|---|---|---|---|
| | iv-RoCi-1.4$^{mRFP}$ | *argB2; pyrG89; veA1; nkuAΔ; IS1::(*A-*An*-P*gpdA-mRFP-An_TtrpC-B*)$_n$ | This study |
| | iv-RoCi-1.5$^{mRFP}$ | *argB2; pyrG89; veA1; nkuAΔ; IS1::(*A-*An*-P*gpdA-mRFP-An_TtrpC-B*)$_n$ | This study |
| | iv-RoCi-2.1$^{mRFP}$ | *argB2; pyrG89; veA1; nkuAΔ; IS1::(*A-*An*-P*gpdA-mRFP-An_TtrpC-B*)$_n$ | This study |
| | iv-RoCi-2.2$^{mRFP}$ | *argB2; pyrG89; veA1; nkuAΔ; IS1::(*A-*An*-P*gpdA-mRFP-An_TtrpC-B*)$_n$ | This study |
| | iv-RoCi-2.3$^{mRFP}$ | *argB2; pyrG89; veA1; nkuAΔ; IS1::(*A-*An*-P*gpdA-mRFP-An_TtrpC-B*)$_n$ | This study |
| | iv-RoCi-2.4$^{mRFP}$ | *argB2; pyrG89; veA1; nkuAΔ; IS1::(*A-*An*-P*gpdA-mRFP-An_TtrpC-B*)$_n$ | This study |
| | iv-RoCi-2.5$^{mRFP}$ | *argB2; pyrG89; veA1; nkuAΔ; IS1::(*A-*An*-P*gpdA-mRFP-An_TtrpC-B*)$_n$ | This study |
| sDIV0525 | iv-RoCi-3.1$^{mRFP}$ | *argB2; pyrG89; veA1; nkuAΔ; IS1::(*A-*An*-P*gpdA-mRFP-An_TtrpC-B*)$_{68}$ | This study |
| | iv-RoCi-3.2$^{mRFP}$ | *argB2; pyrG89; veA1; nkuAΔ; IS1::(*A-*An*-P*gpdA-mRFP-An_TtrpC-B*)$_n$ | This study |
| | iv-RoCi-3.3$^{mRFP}$ | *argB2; pyrG89; veA1; nkuAΔ; IS1::(*A-*An*-P*gpdA-mRFP-An_TtrpC-B*)$_n$ | This study |
| | iv-RoCi-3.4$^{mRFP}$ | *argB2; pyrG89; veA1; nkuAΔ; IS1::(*A-*An*-P*gpdA-mRFP-An_TtrpC-B*)$_n$ | This study |
| | iv-RoCi-3.5$^{mRFP}$ | *argB2; pyrG89; veA1; nkuAΔ; IS1::(*A-*An*-P*gpdA-mRFP-An_TtrpC-B*)$_n$ | This study |
| sDIV0526 | iv-RoCi$^{mRFP}$ | *argB2; pyrG89; veA1; nkuAΔ; IS1::(*A-*An*_P*gpdA-mRFP-An_TtrpC-B)*$_{23}$ | This study |
| sDIV0527 | pyrG-d::mRFP | *argB2; pyrG89; veA1; nkuAΔ; IS1::(*A-*Af_pyrGd-An*_P*gpdA-mRFP-An_TtrpC-B)*$_{15}$ | This study |
| sDIV0528 | SCI$^{cord}$ | *argB2; pyrG89; veA1; nkuAΔ; IS1::*A-*An*_P*tef-An_cns*$_1$*(EAAAK)*$_4$*An_cns*$_2$*-An*_T*tef*-B | This study |
| sDIV0529 | RoCi-1$^{cord}$ | *argB2; pyrG89; veA1; nkuAΔ; IS1::(*A-*An*_P*tef-An_cns*$_1$*(EAAAK)*$_4$*An_cns*$_2$*-An*_T*tef*-B*)n* | This study |

| | | | |
|---|---|---|---|
| sDIV0530 | RoCi-2$^{cord}$ | *argB2; pyrG89; veA1; nkuAΔ; IS1::(*A-*An_*P*tef-An_cns$_1$(EAAAK)$_4$An_cns$_2$-An_*T*tef*-B*)n* | This study |
| sDIV0531 | *COSI-1$^{mCit}$* | *argB2; pyrG89; veA1; nkuAΔ; IS1::*A-*An_*P*gpdA-mCit-An_TtrpC-B* | This study |
| sDIV0532 | *uidA-mRFP 1* | *argB2; pyrG89; veA1; nkuAΔ; IS1::*A-*An_*P*gpdA-uidA-An_TtrpC-B* | This study |
| sDIV0533 | *uidA-mRFP 2* | *argB2; pyrG89; veA1; nkuAΔ; IS1::(*A-*An_*P*gpdA-uidA-An_TtrpC-B-Af_PgpdA-mRFP-An_Ttef)$_6$* - A-*An_*P*gpdA-uidA-An_TtrpC-B* | This study |
| sDIV0534 | *uidA-mRFP 3* | *argB2; pyrG89; veA1; nkuAΔ; IS1::(*A-*An_*P*gpdA-uidA-An_TtrpC-B-Af_PgpdA-mRFP-An_Ttef)$_7$* - A-*An_*P*gpdA-uidA-An_TtrpC-B* | This study |
| sDIV0535 | *uidA-mRFP 4* | *argB2; pyrG89; veA1; nkuAΔ; IS1::(*A-*An_*P*gpdA-uidA-An_TtrpC-B-Af_PgpdA-mRFP-An_Ttef)$_{11}$* - A-*An_*P*gpdA-uidA-An_TtrpC-B* | This study |
| sDIV0536 | *uidA-mRFP 5* | *argB2; pyrG89; veA1; nkuAΔ; IS1::(*A-*An_*P*gpdA-uidA-An_TtrpC-B-Af_PgpdA-mRFP-An_Ttef)$_{14}$* - A-*An_*P*gpdA-uidA-An_TtrpC-B* | This study |
| sDIV0537 | AoSCI-1$^{mRFP}$ | Δ*pyrG,* Δ*ku70, IS1::A-An-PgpdA-mRFP-An_TtrpC-B* | This study |
| sDIV0538 | AoRoCi-1$^{mRFP}$ | Δ*pyrG,* Δ*ku70, IS1::(A-An-PgpdA-mRFP-An_TtrpC-B-Backbone)$_2$* | This study |
| sDIV0539 | AoRoCi-2$^{mRFP}$ | Δ*pyrG,* Δ*ku70, IS1::(A-An-PgpdA-mRFP-An_TtrpC-B-Backbone)$_7$* | This study |
| sDIV0540 | AsnSCI$^{mRFP}$ | *pyrG1, kusAΔ, IS1::A-An-PgpdA-mRFP-An_TtrpC-B* | This study |
| sDIV0541 | AsnRoCi-1$^{mRFP}$ | *pyrG1, kusAΔ, IS1::(A-An-PgpdA-mRFP-An_TtrpC-B-Backbone)$_2$* | This study |
| sDIV0542 | AsnRoCi-2$^{mRFP}$ | *pyrG1, kusAΔ, IS1::(A-An-PgpdA-mRFP-An_TtrpC-B-Backbone)$_7$* | This study |
| sDIV0543 | RoCi$^{mCit}$ | *argB2; pyrG89; veA1; nkuAΔ; IS1::(*A-*An_*P*gpdA-mCit-An_TtrpC-B-Backbone)$_n$* | This study |
| sDIV0544 | RoCi$^{mRFP+mCit}$ | *argB2; pyrG89; veA1; nkuAΔ; IS1::(*A-*An_*P*gpdA-mRFP-An_TtrpC-B-Backbone)$_n$* - (A-*An_*P*gpdA-mCit-An_TtrpC-B-Backbone)$_n$* | This study |
| [11]An- *Aspergillus nidulans*; Asn- *Aspergillus niger*; Ao- *Aspergillus oryzae* | | | |

**Table S3:** Primers used in this study.

| Primer ID | Sequence | Description |
|---|---|---|
| **Primers to amplify linear Gene Targeting Substrates and fragments to generate circular Gene Targeting Substrates by *in vivo* assembly.** | | |
| PR_DIV0432 | AGGTGTAAAAGTAGGGAGCGGTAG | A-FW |
| PR_DIV0435 | GAGGAGAGTGGATGGATAGTCTGG | B-RV |
| PR_DIV3161 | ATGGCCTCCTCCGAGGAC | mRFp FW |
| PR_DIV3162 | TTAGGCGCCGGTGGAGT | mRFP RV |
| PR_DIV3875 | AAAGTCGCTGAGGAGTCTCCTCTTATGTGGTT ATATACGACAACGCTCAGGTGTAAAAGTAGGGA GCGGTAG | A-FW 50 bp |
| PR_DIV3876 | GAGCGTTGTCGTATATAACCACATAAGAGAGGAG ACTCCTCAGCGACTTTGAGGAGAGTGGATGGATA GTCTGG | B-RV 50 bp |
| PR_DIV3927 | ATGTCAUCTCCTACCAGTGCC | USER_Linker-Cns2_FW |
| PR_DIV3928 | ATCGAATGUCCGCTCAATGCTGACTACGACTGAG A | USER_tTEF-Cns2_RV |
| PR_DIV1399 | GGATCCACTTAACGTTACTGAAATC | TtrpC-FW |
| PR_DIV0309 | GGTCTTAAUCGCTTACACAGTACACGAGG | TtrpC-RV |
| PR_DIV3963 | GAGCGTTGTCGTATATAACCACATAAGAGAGGAG ACTCCTCAGCGACTTTGTATTGGGATGAATTTTGT ATGCACGC | Ttef-RV 50 bp |
| **Primers for *mRFP* and *mCit* amplification.** | | |
| PR_DIV3161 | ATGGCCTCCTCCGAGGAC | *mRFP* FW |
| PR_DIV3162 | TTAGGCGCCGGTGGAGT | *mRFP* RV |
| PR_DIV3163 | ATGGTGAGCAAGGGCGAG | *mCit* FW |
| PR_DIV3164 | TTACTTGTACAGCTCGTCCATGC | *mCit* RV |
| **Primers for pDIV1051 construction.** | | |
| PR_DIV3923 | GGGTTTAAUATACCCCGCATATAACCCTCCA | US_PacI-TpyrG*_FW |
| PR_DIV3921 | AGTGGGGAUGCCTCAATTGT | US_pyrG-PgpdA_RV |

| | | |
|---|---|---|
| PR_DIV3922 | ATCCCCACUATTCCCTTGTATCTCTACACACAGG | US_pyrG-PgpdA_FW |
| PR_DIV0660 | GGTCTTAAUACACGAGGACTTCTAGAAAGAAGG A | TtrpC-PacI-RV |
| **Primers for pDIV1050 construction.** | | |
| PR_DIV0312 | GGGTTTAAUCGAGACAGCAGAATCACCG | Ptef-FU-PacI Up |
| PR_DIV3924 | ATAGTCAUGGTGAAGGTTGTGTTATGTTTTG | USER_pTEF-Cns1_RV |
| PR_DIV3925 | ATGACTAUAAACGCATATCTGTCCACT | USER_pTEF-Cns1_FW |
| PR_DIV3926 | ATGACAUCTTTGCTGCCGCTTCCTTTGCTGCTGCT TCCTTAGCCGCTGCTTCCTTTGCAGCCGCCTCGTA TATGCCCACCCTGGATCC | USER_(EAAAK)4-Cns1_RV |
| PR_DIV3927 | ATGTCAUCTCCTACCAGTGCC | USER_Linker-Cns2_FW |
| PR_DIV3928 | ATCGAATGUCCGCTCAATGCTGACTACGACTGAG A | USER_tTEF-Cns2_RV |
| PR_DIV3929 | ACATTCGAUTTATGCCGTTATGACT | USER_tTEF-Cns2_FW |
| PR_DIV0022 | GGTCTTAAUGTATTGGGATGAATTTTGTATGC | Ttef-PacI-RV |
| **Primers for pDIV0941 construction.** | | |
| PR_DIV0310 | GGGTTTAAUATTCCCTTGTATCTCTACACACAGG | PgpdA 2.3-FU-PacI Up |
| PR_DIV3954 | AGGAGAGUGGATGGATAGTCTGG | US_B-RV/A.fumPgpdA |
| PR_DIV3964 | ACTCTCCUCCTTACATCATCTGGTATCTACGCAA GC | US_AfumPgpdA-FW/B |
| PR_DIV3965 | AGGCCAUTGTGTAGATTCGTCTGGTACTGAG | US_AfumPgpdA-RV/mRFP |
| PR_DIV3966 | ATGGCCUCCTCCGAGGAC | US_mRFP-FW |
| PR_DIV0022 | GGTCTTAAUGTATTGGGATGAATTTTGTATGC | Ttef-PacI-RV |
| **Primers for pDIV1049 construction.** | | |
| CSN438 | GGGTTTAAUGATCACATAGATGCTCGGTTGACA | U3 promoter Fw |
| PR_DIV4046 | ACCCCCAUCGGCGTGCATCATCCGTGAATCGAA | mCit PS I RV |

| | | |
|---|---|---|
| PR_DIV4045 | ATGGGGGUGTTCTGCGTTTTAGAGCTAGAAATAGC | mCit PS I FW |
| CSN790 | GGTCTTAAUACCCTGAGAAGATAGATGTGAATGTG | U3 terminator Rv |

**Table S4:** Protospacers used in this study.

| **Protospacer ID** | **Sequence** | **Used for** |
|---|---|---|
| *uidA-gRNA* | CGCAGGTGATCGGACGCGTC | *uidA* DSB |
| *mCit-gRNA* | CGCCGATGGGGGTGTTCTGC | *mCit* DSB |

**Table S5**: Probes for ddPCR used in this study.

| **Probe ID**[1] | **Sequence** | **Type** |
|---|---|---|
| Probe_*mRFP* | CCCGTAATGCAGAAGAAGACCATGGG | FAM-Zen quencher-Iowa Black |
| Probe_An_*oliC*-HEX | ACCAAGACCGATGGCAGCAGAG | HEX-Zen quencher-Iowa Black |
| Probe_Asn_*oliC*-HEX | AACCTGGGTATGGGTTCCGCTG | HEX-Zen quencher-Iowa Black |
| Probe_Ao_*oliC*-HEX | ACCGAAGACGAGACCGATACCGAT | HEX-Zen quencher-Iowa Black |

[1]An- *Aspergillus nidulans*; Asn- *Aspergillus niger*; Ao- *Aspergillus oryzae*

**Table S6.** Copy numbers of the top five most fluorescent strains of three transformations made with a pre-formed c-GTS (RoCi) and of three transformations made with a c-GTS made by in vivo assembly (iv-RoCi) ordered from highest to lowest in copy-number.

| **RoCi 1** | **RoCi 2** | **RoCi 3** | **iv-RoCi 1** | **iv-RoCi 2** | **iv-RoCi 3** |
|---|---|---|---|---|---|
| 5,16 ± 0,44 | 10,3 ± 1 | 17,5 ± 0,8 | 22,7 ± 2,1 | 26,8 ± 1,25 | 68 ± 6 |
| 4,47 ± 0,54 | 7,07 ± 0,23 | 7 ± 1,3 | 21,1 ± 1,3 | 25,5 ± 6,92 | 32,5 ± 10,4 |
| 4,27 ± 0,2 | 4,18 ± 0,32 | 5,48 ± 0,36 | 20,2 ± 1,1 | 22,7 ± 3,21 | 32 ± 6 |
| 2,43 ± 0,07 | 3,8 ± 0,8 | 3,29 ± 0,13 | 20 ± 1,3 | 12 ± 1,14 | 24,1 ± 2,7 |
| 3,17 ± 0,09 | 3,57 ± 0,26 | 2,63 ± 0,08 | 10,9 ± 2,52 | 5,03 ± 0,19 | 20 ± 1,4 |

### References


1. Hansen BG, Salomonsen B, Nielsen MT, Nielsen JB, Hansen NB, Nielsen KF, et al. Versatile Enzyme Expression and Characterization System for Aspergillus nidulans, with the Penicillium brevicompactum Polyketide Synthase Gene from the Mycophenolic Acid Gene Cluster as a Test Case. *Appl Environ Microbiol*. 2011 May;77(9):3044–51.

2. Nødvig CS, Nielsen JB, Kogle ME, Mortensen UH. A CRISPR-Cas9 System for Genetic Engineering of Filamentous Fungi. *PLoS One*. 2015;10(7):e0133085.

3. Jarczynska ZD, Rendsvig JKH, Pagels N, Viana VR, Nødvig CS, Kirchner FH, et al. DIVERSIFY: A Fungal Multispecies Gene Expression Platform. *ACS Synth Biol*. 2021 Mar 19;10(3):579–88.
